# Mitigating memory effects during undulatory locomotion on hysteretic materials

**DOI:** 10.1101/748186

**Authors:** Perrin E. Schiebel, Henry C. Astley, Jennifer M. Rieser, Shashank Agarwal, Christian Hubicki, Alex M. Hubbard, Kelimar Cruz, Joseph Mendelson, Ken Kamrin, Daniel I. Goldman

## Abstract

Undulatory swimming in flowing media like water is well-studied, but little is known about locomotion in environments that are permanently deformed by body–substrate interactions like snakes in sand, eels in mud, and nematode worms in rotting fruit. We study the desert-specialist snake *Chionactis occipitalis* traversing granular matter and find body inertia is negligible despite rapid transit and speed dependent granular reaction forces. New surface resistive force theory (RFT) calculation reveals how this snakes wave shape minimizes memory effects and optimizes escape performance given physiological limitations (power). RFT explains the morphology and waveform dependent performance of a diversity of non-sand-specialist, but overpredicts the capability of snakes with high slip. Robophysical experiments recapitulate aspects of these failure-prone snakes and elucidate how reencountering previously remodeled material hinders performance. This study reveals how memory effects stymied the locomotion of a diversity of snakes in our previous studies [Marvi et al, Science, 2014] and suggests the existence of a predictive model for history-dependent granular physics.

## I. INTRODUCTION

Movement is critical to the survival of many organisms and a necessary ability in robots used in fields like medicine [1], search and rescue [2], and extraterrestrial exploration [3]. This complex phenomenon emerges from the interplay between a locomotor’s internal body shape changes and the physics of the surroundings. Successful locomotion thus depends on the execution of self-deformations which generate appropriate reaction forces from the terrain; a relationship which can be further complicated if the body motion permanently changes the state of the substrate. Much of our knowledge of terrestrial locomotion is in the regime of rigid materials where the terrain is not affected by passage of the animal [4–7], leading to the development of robots which are effective on hard ground [8–10].

Little is known about locomotion in non-rigid materials which are plastically deformed by the movement of animals or robots, leaving tracks or footprints. Deformable terrains inhabit a spectrum from materials like water which continuously flow toward the undisturbed, zero shear state [11] to those which are remodeled by the interaction, like sand. Most work on motion in yielding materials has been on systems in which disturbances dissipate (fluids) [12–14]; the impact of soft material hysteresis on locomotion is not well-understood [15].

At one end of the spectrum, where deformations are short-lived [16], small fluid swimmers like the nematode *Caenorhabditis elegans* [17], spermatazoa [18], and bacteria in water [19] and macroscale frictional-fluid swimmers like the sandfish lizard and shovel-nosed snake moving subsurface through sand [5] use the resistance of the surrounding material to the motion of their body shape changes to propel themselves. The fluid in these non-inertial systems continuously re-flows around the body of the animal such that the state of the material is largely unchanged by the swimmer on time-scales relevant to locomotion.

At the other end of the spectrum is movement on the surface of materials with memory [22], like sand. These substrates flow in response to stress, but, unlike fluids, they remain in the altered state as exemplified by the tracks left behind by a passing animal. We previously studied the crutching motion of mudskippers and turtles which use limbs to traverse granular matter (GM) substrates. These animals managed the changes they introduced to the material by avoiding their own tracks with large-enough steps [23, 24]. Sidewinder rattlesnakes employ a similar strategy, using a specialized two-wave waveform to “step” across the surface, minimizing slipping and avoiding their own tracks by creating static contacts with the substrate [25, 26].

Limbless, undulatory animals like snakes are uniquely coupled to their surroundings. When moving via *lateral undulation*, the slithering gait typically associated with elongate, limbless vertebrates [25], the body is continuously experiencing drag as it slides through the terrain. These organisms can generate normal forces to counteract body drag by pushing the sides of the trunk against heterogeneity in the surroundings [27]. How do such slithering animals, which cannot take larger steps to avoid their previously-laid tracks, move effectively on material where their interactions permanently deform the substrate?

This is a challenging task as demonstrated by the failure of both a snake-like robot [28] and a number of snake species [26] attempting to traverse the surface of GM. Better understanding of the connection between body self-deformations, substrate remodeling and force generation, and the resulting locomotor performance will elucidate principles for effective motion on hysteretic materials. Future studies can leverage such knowledge of the benefits and limitations of different body–terrain interaction modes to tease apart neuromechanical control strategies for contending with natural terrains [29].

In this paper we studied this system using animal experiments, granular drag measurements, RFT calculations, and a robophysical model. In Section II we quantified the kinematics of snakes traversing a GM substrate. In Sec. II A we compared a variety of species with a sand-specialist. We measured the granular response to surface drag in Sec. III. In Sec. III A we developed a model for speed dependence and in Sec. III B found a characteristic drag anisotropy curve. The speed and depth independent anisotropy motivated the use of RFT in Section IV to calculate the relationship between waveform and performance. In Sec. IV A we identified a trade off between actuation speed and torque from the GM which we show in Sec. IV B explained the stereotyped shape used by the sand-specialist. We show in Sec. IV C that performance of the snakes depended on both morphology and waveform. In Section V we systematically explored the connection between waveform, substrate, and performance using a robophysical model (Sec. V A). In Sec. V B we applied RFT to the robot. We found in Sec. V C RFT was inaccurate because the robot was re-interacting with remodeled material. RFT could provide insight into the regime of locomotor failure in Section V D, where we found the sand-specialist remains far from failure due to a combination of its waveform, slender body, low-friction scales, and lifting of segments that only produce drag. We close in Section VI with a summary of our results and a brief discussion of the implications.

## II. PERFORMANCE OF LIMBLESS LOCOMOTORS ON GRANULAR MATERIAL

Snakes occupy a variety of habitats and display a wide range of naturally occurring morphologies. We took advantage of this natural diversity to explore how different body shapes and patterns of self-deformation fare on GM. We studied 22 species of snakes representing five families from the collection at Zoo Atlanta. The snakes’ body plans ranged from short and stout to long and slender (Fig. 1(a,b)) and their natural habitats encompassed a broad range from wetlands and swamps to wet and dry forests and rainforest canopies to deserts to rocky mountains.

**FIG. 1.**
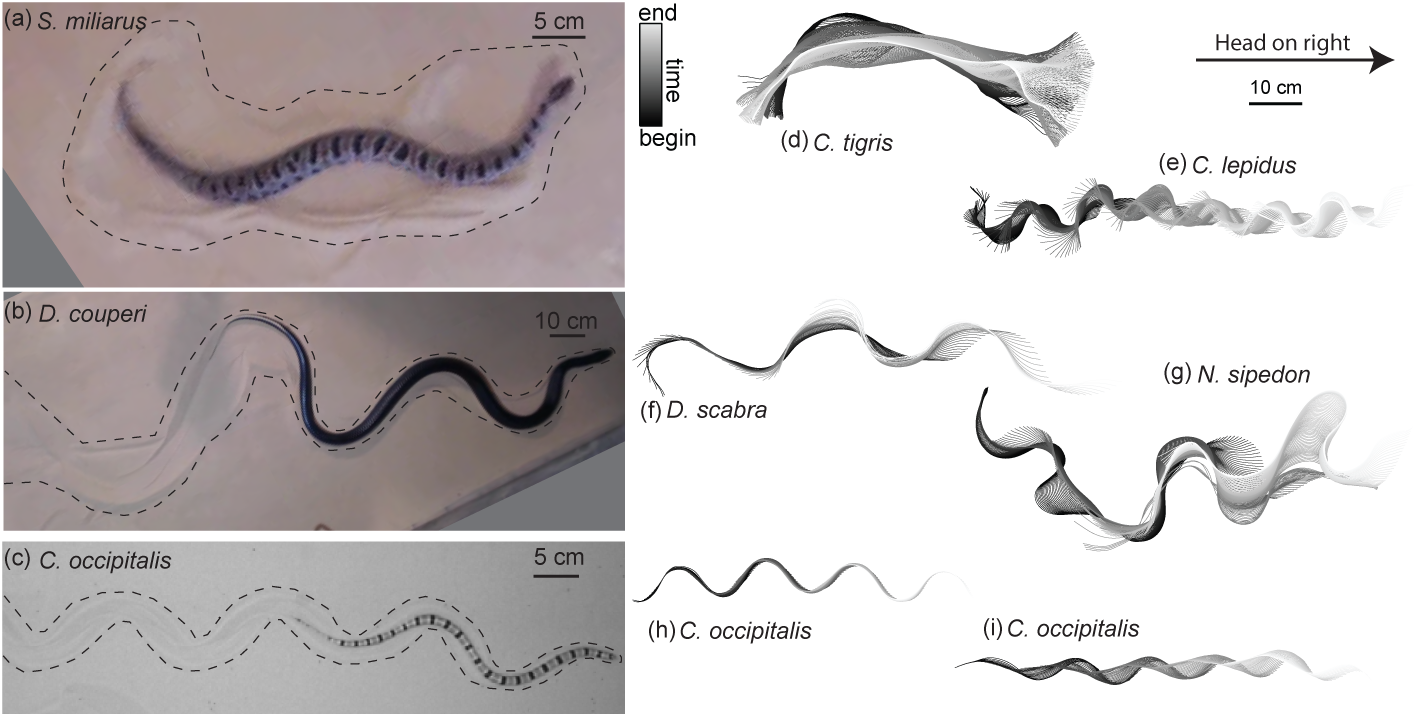
Body shape, waveform, and ability to progress across GM varies among snake species. (a-c)Snapshots of snakes moving on the surface of GM. Dashed line roughly indicates the area of material disturbed by the motion of the animal. (a) The generalist pygmy rattlesnake *Sistrurus miliarius*, which lives throughout the southeastern US, attempting to move on natural sand collected from Yuma, Arizona, USA. The animal has completed several undulations, sweeping GM lateral to the midline of the body. This snake failed to progress further than pictured. (b) The generalist eastern indigo snake *Drymarchon couperi* lives in the southeastern US, although its range is smaller than *S. miliarus*, primarily inhabiting Florida. On the same GM as (a) (c) The sand-specialist shovel-nosed snake *Chionactis occipitalis* in the lab on 300 *µ*m glass particles. (d-i) Digitized midlines of animals. Color indicates time from beginning to end of the trial. All scaled to the 10 cm scale bar shown. (d-g) are on the Yuma sand and (h,i) are on glass particles. (d) *Crotalus tigris*. This snake superficially shares some of its natural range with that of *C. occipitalis*, but is closely associated with rocky boulder substrates rather than sand. The animal was unable to progress on the GM. Total length of the trial *t*_*tot*_ =25.7 s, time between plotted midlines *δt* = 100 ms (e) *Crotalus lepidus*, a generalist in the southwestern US and the central part of Mexico. *t*_*tot*_ =9.4 s, *δt* = 33 ms (f) *Dasypeltis scabra* inhabits a wide range of habitats in Africa. *t*_*tot*_ =1.6 s, *δt* = 33 ms (g) *Nerodia sipedon*, a water snake inhabiting most of the Eastern US extending into Canada. *t*_*tot*_ =6.0 s, *δt* = 33 ms (h) *C. occipitalis* 130 and (i) 128. These trials represent the intra-individual variation in kinematics. Some of the animals moved approximately “in a tube” where all segments of the body followed in the path of their rostral neighbors (e.g. (h), *t*_*tot*_ =1.25 s, *δt* = 12 ms) while others used this strategy only on the anterior portion of the body, appearing to drag the posterior segments in a more or less straight line behind themselves (e.g. (i), *t*_*tot*_ =1.01 s, *δt* = 12 ms). See [21] for tables of length, width, and mass of all animals used.

As a counterpoint to the variety of snakes, which were either terrain generalists or specialized to habitats which did not have an omnipresent granular substrate, we also studied the shovel-nosed snake *Chionactis occipitalis* (Fig. 1(c)). This species is specialized to bury within [5, 30] and move across [31] the dry sand of their desert habitat. We used nine individuals collected from the desert in Arizona, USA (Appendix B). These sand-specialists are a consummate example of a successful strategy for moving across material with memory; they use a stereotyped waveform ([21], [29]) to quickly traverse many body lengths across the sand with little slipping of the body, leaving behind distinct tracks [31].

We focused our study on snakes using lateral undulation. This gait is characterized by flexions of the trunk, primarily in the plane parallel to the substrate, which are passed from head to tail. If heterogeneity in the surroundings resists tailward slipping of these bends, the result is that the bends are stationary with respect to the terrain and the snake’s body is propelled forward [25].

### A. Kinematics and ability vary among species

In the laboratory we studied the non-sand-specialist species moving on a trackway of natural sand (Appendix A, videos collected for [26]) and nine *C. occipitalis* individuals on a previously-used model for natural sand, ≈ 300 *µ*m glass particles ([5, 32], Appendix B). Overhead high-speed cameras captured the trials and the GM was prepared to an undisturbed initial state using a fluidized trackway (Appendix A,B).

The use of alternating left and right bends on the body was ubiquitous, although specifics of the waveform such as amplitude and wavenumber varied (Fig. 1). Among the non-specialists some snakes used uniform, periodically repeating bends (e.g. Fig. 1(b,e,f)) like *C. occipitalis* (Fig. 1(c,h,i). Others used curves of varying amplitude and wavelength along the body (Fig. 1(g)).

We digitized the midline of the animals and used the tangent angle, *θ* (Fig. 2(a),*left*), to represent body posture at each instant in time. We characterized the waveform of each snake by measuring the maximum tangent angle, *θ*_*m*_, also referred to as the attack angle, and the number of waves on the body, *ξ* (Fig. 2(a),*right*, Appendix A,B). For an infinitesimally thin curve with infinite resolution these parameters are independent. In practice, the musculoskeletal system and width of the body limits both how quickly the midline can change curvature and how sharply bent the body can be, although sufficiently far away from these limits *θ*_*m*_ and *ξ* remain independent. While there may be other, not purely anatomical, factors which connect these variables, in this work we chose to measure their values and study the connection to performance as mediated by the GM. Future study could explore the neuromechanical origin of the shapes and its connection to the differences observed between different species.

**FIG. 2.**
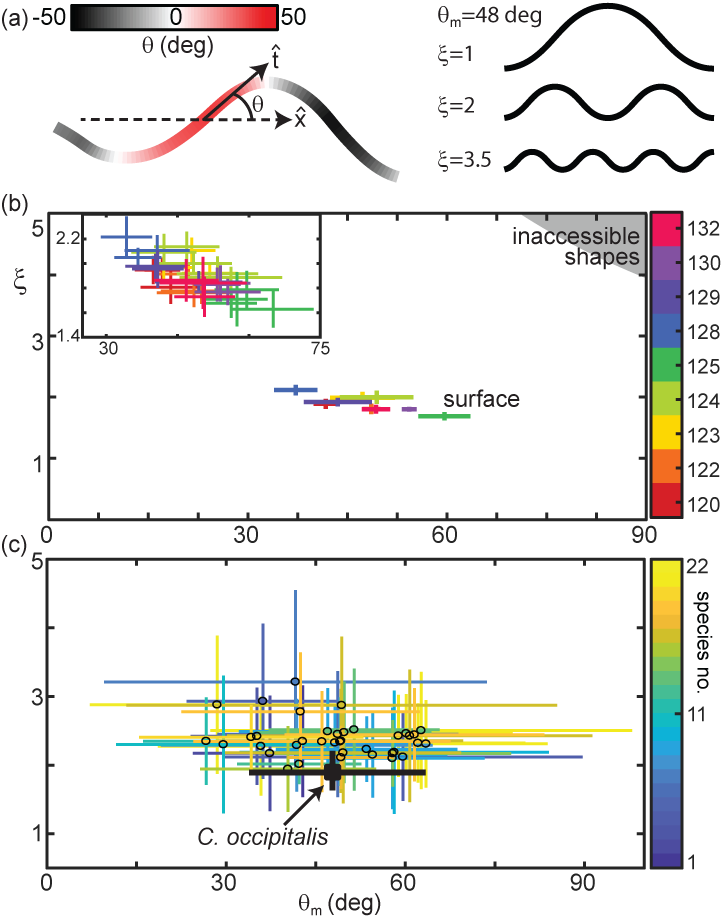
Snake waveform parameters measured in experiment. (a, *left*) Tangent angle, *θ* is the angle between 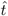 and the average direction of motion of the animal, 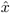. Example body posture shown, colored by *θ*. (a, *right*) *ξ* is the spatial frequency. Shown are example serpenoid (sinusoidal tangent angle) curves on a body of fixed length and *θ*_*m*_ = 48 deg. Shown are *ξ* = 1, 2, and 3.5 waves on the body. (b) *C. occipitalis* experimental measurements plotted in the (*θ*_*m*_, *ξ*) parameter space. Color indicates animal number. Markers are the mean and range of each individual taken over all trials. N=9 individuals, n=30 trials. The gray region in the upper right corner are waves which are inaccessible given the flexibility of the snake [5]. *θ*_*m*_ was comparable to values measured from images of tracks taken in the field [21]. (*inset*) shows a close-up of the data with mean and range measured in each trial plotted individually and color consistent with the main plot (Linear fit with 95% confidence interval to inset data: slope −0.016 (−0.020,-0.011) intercept 2.66 (2.42, 2.90), R-square=0.6). (c) Axes are as in (b), black cross represents the average *C. occipitalis* measurements. Colored crosses are the mean and range of values measured on a per-trial basis using the various non-specialist species, indicated by color. N=22, n=38.

We measured *θ*_*m*_ and *ξ* for each of the nine *C. occipitalis* and found that the animals used a limited subset of the range of the shapes they were anatomically capable of adopting (Fig. 2(b), colored crosses). There was a slight downward trend in the relationship between *ξ* and *θ*_*m*_ when considering both each individual (Fig. 2(b)) and each trial (Fig. 2(b), *inset*). Given the weak relationship between *ξ* and *θ*_*m*_, we decided to combine all of the individual measurements and explore the impact of the average *C. occipitalis* wave, taken as the mean and range of the averages shown (Fig. 2(c), black cross).

Compared to *C. occipitalis*, there was greater variety among the wave parameters measured on the non-specialist species (Fig. 2(c)). However, as in *C. occipitalis*, the parameters did not fill the space of anatomically possible shapes, suggesting the animals were operating under some constraints. To better understand what factors contributed to the choice of waveform we studied the granular response to drag to characterize the forces experienced by the animals.

## III. GRANULAR DRAG MEASUREMENTS

The forces acting on animals moving submerged within GM were elucidated by subsurface drag measurements [32]. These experiments revealed that during subsurface swimming in GM the material acts like a frictional fluid; propulsive forces arise from the resistance of the GM to motion of the body segments as the grains interact with each other via normal and frictional contacts [15, 32]. A similar understanding of the character of movement at the surface is unknown.

We observed the formation of granular piles as the snakes self-deformed on the granular surface (Fig. 3(a), [21]). This suggested that, consistent with subsurface swimming and surface walking, slithering animals propel themselves using the granular forces arising from the body yielding the material (and not solely frictional anisotropy of the ventral scutes [33]). The interaction between the body and the GM can be characterized by the amount of slip, 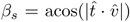, relative to the substrate, where 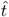 and 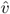 are the local velocity and tangent unit vectors of a body segment (Fig. 3(a)*inset* [5]). *β*_*s*_ emerges from the interrelationship between the granular stress and the animal’s self-deformation pattern.

**FIG. 3.**
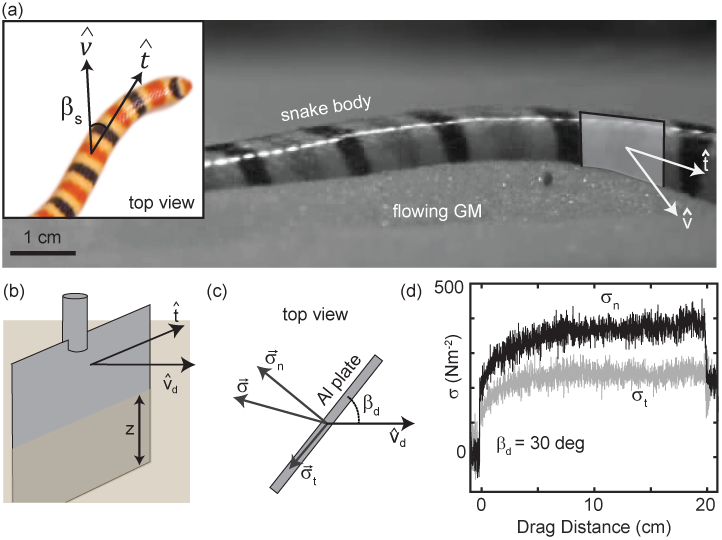
Characterizing granular forces experienced by snakes by empirically measuring stress on a partially buried plate. (a) Side-view of *C. occipitalis* moving on the surface. Snake is moving from left to right. The surface was initially featureless, the piles of sand were generated by the motion of the snake [21]. We chose the simplest model of a snake body segment, a flat plate aligned with 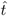 of the midline moving in direction 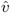. 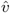 is an amalgamation of the segment movement associated with the self-deformation and the center-of-mass (CoM) speed arising from interaction with the GM. (*inset*) Top view of a snake. *β*_*s*_ is the angle between the local tangent and velocity unit vectors. (b) Aluminum plate model of a snake segment, dimensions 3 × 1.5 × 0.3 cm^3^. The plate was kept at a constant depth, *z*, from the undisturbed free surface of the GM to the bottom of the intruder. (c) Top view of the plate moving in a direction 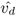 at angle *β*_*d*_ between the direction of motion and the tangent to the plate face. A force transducer decomposed the total stress, *σ* into *σ*_*t*_ and *σ*_*n*_. (d) Raw drag data collected at *β*_*d*_ = 30° and *v*_*d*_ = 10 mm s^−1^ as a function of drag distance of the plate. The upper black curve is *σ*_*n*_, the gray is *σ*_*t*_. Note plotted data is down sampled by a factor of 10.

We empirically measured the granular stress on a simple model for a snake body segment—an aluminum plate commanded to move at a slip angle *β*_*d*_ between the drag velocity unit vector 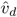 and the plate face tangent in the horizontal plane (Fig. 3(a,b)). We dragged the plate for 20 cm at a constant depth, *z*, measured from the intruder’s bottom edge (Fig. 3(b)) and measured stress normal, *σ*_*n*_, and tangent, *σ*_*t*_, to the plate face (Fig. 3(c), Appendix E). A fluidizing bed containing the same 297 ± 40 *µ*m glass particles used in the *C. occipitalis* experiments prepared the material to an initially loose-packed state.

Unlike subsurface drag stresses, which developed almost instantaneously to the steady state [32], at the surface stress monotonically increased over several centimeters before saturating (Fig. 3(d)). This is due to the free surface flow of the GM; a pile of sand above the surface, like those created by the snakes, appeared at the leading face of the intruder at the onset of drag and increased in volume until reaching a balance between the new grains being encountered and those flowing around the edges of the plate. Previous studies of plate drag at the surface (at *β*_*d*_ = 90 deg) measured a similar drag distance dependent force and observed a wedge-shaped region of GM, beginning at the bottom edge of the plate and extending above the surface, whose constituent grains were flowing forward and up against gravity [34, 35].

### A. Velocity-dependent stress captured by grain inertia model

The snake speeds were variable (segment speeds from ∼ 35 to 95 cm s^−1^ [21]) and the intrusion depth of the snakes’ trunk into the GM ranged from 0 (no intrusion, occurring at the apexes of the wave which the snake lifted off of the surface) to ∼ 5 mm [21]. Previous studies indicated that granular drag stress depends on both intruder speed [36] and depth [26, 37]. Commensurate with those studies, we fixed *β*_*d*_ = 90 deg and *z* = 8 mm and found normal stress quadratically increased as *v*_*d*_ increased from 1 mm s^−1^ to 750 mm s^−1^ (the limit of the robot arm capability, Fig. 4(a)). Similarly, for *β*_*d*_ = 90 deg and *v*_*d*_ = 10 mm s^−1^, normal stress linearly increased with *z* from 4 mm, the shallowest depth where force could be resolved, to 40 mm, where the plate was fully submerged with the top edge 10 mm below the surface (Fig. 4(b)).

**FIG. 4.**
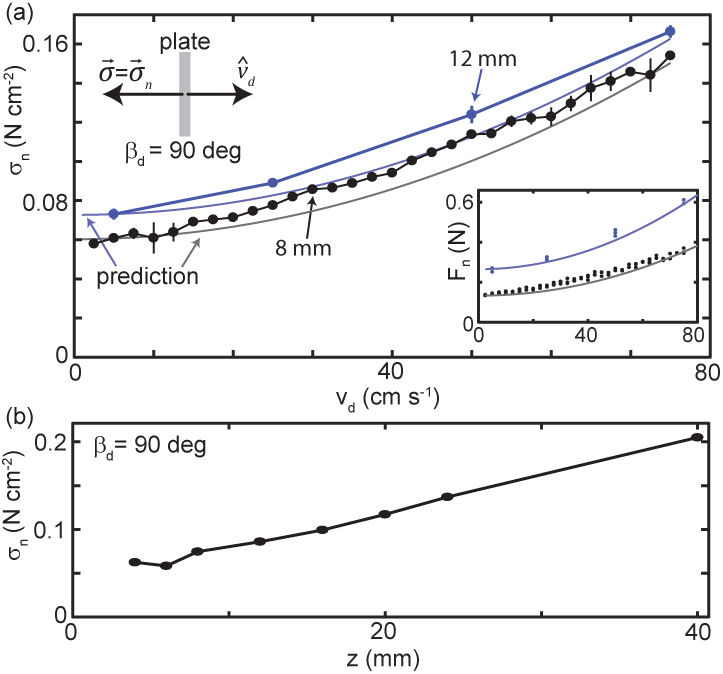
Granular drag stress as a function of speed and depth. (a) Stress normal to the plate face as a function of *v*_*d*_ at a constant depth *z* = 8 mm (black points) and *z* = 12 mm (blue points). *β*_*d*_ = 90° for all trials such that the total stress was equal to *σ*_*n*_. Circle markers are mean and error is std. of three trials. Gray curve is model prediction for *z* = 8 mm and light blue curve for *z* = 12 mm. (*inset*) Normal force versus *v*_*d*_ measured in experiment (circle markers) and as predicted by the model (solid curves). Color is consistent with main plot. Each dot is the average force measured in one trial, all trials shown. (b) Stress normal to the plate face as a function of *z* at *v*_*d*_ = 10 mm s^−1^.

For snakes moving slowly on non-deformable surfaces, body inertia was small compared to the frictional forces between the belly and the surface [33]. While *C. occipitalis* was moving quickly, we observed, in line with earlier reports [38], that forward motion would immediately cease when the animals stopped propagating the wave. This phenomena is observed in swimmers at low-Re like *C. elegans* and bacteria where the dominant resistive forces of the surroundings inhibit gliding.

The animal behavior indicated that friction between the body and the GM and dissipative interactions within the GM had greater impact on the motion than inertia of the body. We thus assumed a resistive-force-dominated system in which stress arose from independent static and dynamic terms.

We modelled stress on the plate as a static stress *σ*_*o*_(*v*_*d*_ → 0), assumed to arise from Coulomb-wedge type forces [35], plus a material inertia term 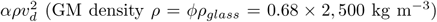. We estimated a packing fraction *ϕ* = 0.58 induced by motion of the intruder with a scalar *α* which was expected to be of *O*(1). We estimated *σ*_*o*_ using the experimental data by subtracting 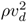 from *σ*_*n*_ measured at the three lowest speeds and averaging the result. Our prediction captured stress as a function of *v*_*d*_ (Fig. 4(a), gray and blue curves) with a pre-factor value of *α* = 1.1. The model tended to under-predict stress, although the bulk of the effect of speed was captured. The predictive power of this simple model, for which the form of the static and dynamic contributions did not change across the range of *v*_*d*_ tested, supported our hypothesis that, across the relevant drag speeds, the material was in the regime where local, resistive forces dominated.

Eels in water (a high-Re system) increase the CoM velocity, *v*_*CoM*_, by both increasing the speed of wave propagation and altering the waveform to manage the speed-dependent fluid response [39]. While the wave parameters of *C. occipitalis* did not systematically vary across the range of speeds tested, the relationship between the speed of wave propagation and *v*_*CoM*_ was linear [21]. This suggested that the aggregate forces experienced by the animal depended only on the pattern of self-deformation and not other factors such as the speed of self-deformation.

### B. Drag anisotropy is not strongly dependent on speed or depth

At low-Re, the normal and tangential stresses acting on segments of long, slender swimmers in Newtonian fluids are approximated by *σ*_*n*_ = *C*_*n*_sin(*β*_*d*_) and *σ*_*t*_ = *C*_*t*_cos(*β*_*d*_), respectively [40]. The constants *C*_*n*_ and *C*_*t*_ are determined by the geometry of a body segment and the viscosity of the surrounding fluid, and the ratio *C*_*n*_*/C*_*t*_ can be used to approximate swimming performance of a given waveform [16]. We assumed that, given body inertia was small compared to forces from the GM resisting motion, the performance of the snake was largely a function of the ratio of propulsive to drag stresses, 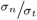 (further supported by RFT calculations in the following Sec. IV).

We measured stress as a function of *β*_*d*_ ranging from 0 deg to 90 deg (Fig. 3(b)). The ratio 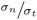 was largely independent of the drag distance (Fig. 5(a)), especially at small *β*_*d*_. There was a periodic fluctuation of 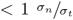 occurring over several cm appearing in all trials. However, both these fluctuations and the slope of 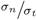 as a function of drag distance were small relative to the effect of changing *β*_*d*_. Thus, we averaged *σ*_*n*_ and *σ*_*t*_ to characterize the relationship between plate orientation and stress normal and tangential to its face (Fig. 5(b)).

**FIG. 5.**
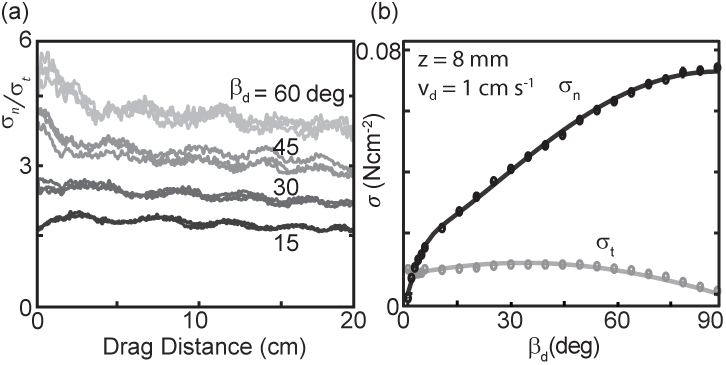
Stress as a function of plate drag angle. (a) Stress anisotropy versus drag distance. Since *σ*_*t*_ is small, values with noise included can approach zero. As we were interested in the force evolution over several cm (for the plotted drag speed this corresponds to signals < 1 Hz), we removed fluctuations above 5 Hz using a low-pass butterworth filter. Curves from three trials are shown for each of four different, fixed *β*_*d*_ = 15 (darkest, bottom curve), 30, 45, and 60 deg (lightest, top curve) as labeled on the right of the plot. The slopes of a linear regression fit to the average of three trials as a function of drag distance were −1.8 × 10^−5^, −2.9 × 10^−5^, −5.0 × 10^−5^, and −6.4 × 10^−5^ cm^−1^ for *β*_*d*_ = 15, 30, 45, and 60°, respectively (R-square=0.6, 0.8, 0.8, and 0.7). (b) Average stress as a function of changing drag angle. Each measurement was calculated by averaging the raw stress data from 10 to 20 cm drag distance. Each data point is the mean and standard deviation of three trials; error bars are smaller than the markers. Solid lines are the fit functions used in the RFT calculations.

We found that normal stress increased monotonically with *β*_*d*_ while tangential stress was approximately constant, gradually falling to zero as *β*_*d*_ went to 90° (Fig. 5(b)). The ratio 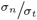 as a function of *β*_*d*_ for both varying speeds and depths collapsed to a characteristic anisotropy curve (Fig. 6(a)). The same drag anisotropy curves were measured in poppy seeds and oolite sand and were similarly independent of depth and speed [24]. The comprehensive appearance of this curve suggests it may be a more general feature of dissipative, deformable materials.

**FIG. 6.**
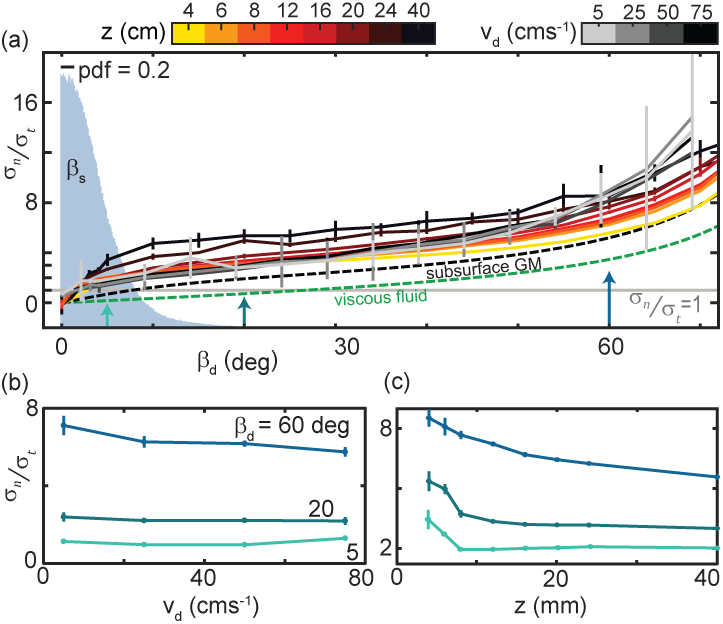
The ratio of granular normal to tangential stress, 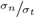, is not strongly dependent on speed or depth. (a) 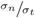 as a function of *β*_*d*_. Light gray horizontal line indicates 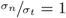. All depths and speeds are plotted as indicated by color. Dashed curves are the anistropy for a low-Re swimmer in Newtonian fluid (green, *K*_*fluid*_ ≈ 1.5 for a smooth, long, and slender swimmer [16]) and subsurface in the same ≈ 300 *µ*m GM used in this study (black). Labels are below the associated curves. The solid gray area is the probability density (pdf) of *β*_*s*_ measured on the snake in experiment (N=9, n=30). (b) 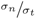 as a function of *v*_*d*_ at the values of *β*_*s*_ indicated. Colors correspond to the colored arrows on the horizontal axis in (a). Linear fits to the data (not shown) with slope, m, and 95% confidence bounds in parentheses for *β*_*d*_ = 60° *m* = −0.019°(− 0.040, 0.003) R-square=0.88; *β*_*d*_ = 20° *m* = − 0.003° (− 0.007, 0.002) R-square=0.75; *β*_*d*_ = 5° *m* = − 0.003° (− 0.011, 0.016) R-square=0.23, (c) 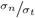 as a function of *z*. Colors and angles are the same as in (b).

Because tangential stress magnitudes were relatively small and normal stresses rose sharply with *β*_*d*_ at small angles (Fig. 5(b)), thrust and drag forces on the plate were equal 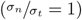 at smaller *β*_*d*_ than in viscous fluid or subsurface in GM (Fig. 6(a)). Reflecting the small angles where the anisotropy was unity, *β*_*s*_ measured on the snake were small (Fig. 6(a), gray histogram).

Plotting anisotropy at constant *β*_*d*_ as speed and depth varied further illustrated the relatively weak dependence of 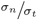 on these variables (Fig. 6(c,d)). Especially at small *β*_*d*_, anisotropy did not depend on *v*_*d*_. 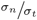 was more dependent on *z* at shallow depths, decreasing twofold as *z* increased from 4 to 8 mm (Fig. 6(c)).

## IV. SURFACE RESISTIVE FORCE THEORY MODEL

Continuum constitutive equations for granular media (similar to Navier-Stokes equations for fluids) can predict granular flows [41]. However, these methods, as well as well established discrete element methods (like MD), are computationally expensive. Granular resistive force theory, which relies on the assumption that the force on the body is a linear, independent superposition of the forces on individual segments [40], is by comparison computationally inexpensive. RFT successfully models a number of systems for which these assumptions are valid, although its effectiveness in a system exhibiting hysteresis was unknown [15].

The negligible body inertia and local, dissipative nature of the forces indicated using resistive force theory (RFT, [40]) and assuming quasi-static locomotion, such that forces were balanced at each moment, was a good candidate for probing the connection between waveform and performance of the surface slithering snakes. The insensitivity of anisotropy on speed suggested that the same speed-independent granular RFT used to predict subsurface performance in GM [15, 32] could be applied to this system.

The empirical relationships between body segment motion, characterized by *β*_*d*_, and stress measured in the drag experiments (Fig. 5(b)) provided the necessary information to estimate the forces acting on different waveforms using RFT.

We first focused on calculation of sand-specialist performance. RFT, using the force relations for *v*_*d*_ = 10 mm s^−1^ and *z* = 8 mm, accurately predicted the average *v*_*CoM*_ and the relationship between *v*_*CoM*_ and *v*_*seg*_ of *C. occipitalis* for the average snake kinematics and morphology [21]. We also performed test RFT calculations using the drag stresses measured at each depth and found that the effect of depth was much less than that of shape and less than the experimental uncertainty [21]. The prediction was similarly insensitive to v_d_ [21].

At the apexes of the body wave body segments experience drag without contributing thrust. Previous research found snakes moving on firm surfaces lifted these sections of the body off of the substrate [33]. In the lab we measured 3D kinematics and found that *C. occipitalis* lifted the wave apexes such that there was a vertical wave on the body which was twice the spatial and temporal frequency of the horizontal wave (Appendix C, [21]). These lifted segments are at *β*_*s*_ near 0 deg, where the drag is greater than the thrust (*K* < 1) (Fig. 6(a)). Therefore, removing these segments from contact with the material reduces drag without a decrease in propulsive force. We included lifting in the RFT calculation by assigning zero force to those segments whose *θ* was in the lowest 41% (equivalent to the wave apexes [21]). While the predicted benefit of lifting to this species was small because of its low-friction scale and a slender body, tracks in the wild also show evidence of lifting.

Subsurface GM swimming performance is improved by reducing scale friction [5]. The surface granular RFT similarly predicted that slithering speed decreased as scale friction increased [21]. This is an intuitive result as increasing the tangential stress shifts the anisotropy curve down, moving the location of 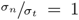 to larger *β*_*d*_ (Fig. 6(a)).

### Trade off between internal and external factors constrains waveforms

We used RFT to predict the maximum joint torque, *τ*_*m,RFT*_, experienced by any *C. occipitalis* body segment over an undulation cycle (assuming infinitesimal width and finite segment length, Fig. 7(a), Appendix F). *τ*_*m,RFT*_ increased as *θ*_*m*_ or *ξ* decreased both because balancing forces required larger *β*_*d*_, thus greater force magnitudes (Fig. 5(b)), and these shapes “stretched out” the body, creating longer lever arms (see Fig. 7(b) and compare shapes in lower left corner to upper right).

**FIG. 7.**
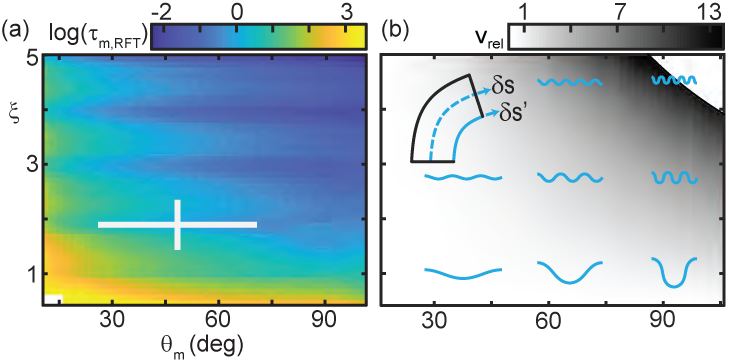
Trade-off between external torque and internal actuation speed. (a) RFT prediction of the peak joint level torque, *τ*_*m,RFT*_, occurring during one undulation cycle as a function of *θ*_*m*_ and *ξ*. Color indicates the log of *τ*_*m,RFT*_. White cross is the mean and range measured in experiment. (b) Heatmap of relative actuation speed,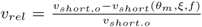. *v*_*rel*_ = 1 at the nominal waveform of *C. occipitalis* (*θ*_*m*_ = 48.4, *ξ* = 1.9). Frequency, *f*, is calculated to yield the desired *v*_*CoM*_for a given waveform’s stride length (Appendix G). Diagram in upper left corner illustrates the bending beam. The nominal length of a segment at the midline is *δs*, blue dashed line. The length of the inside of a segment is *δs′*, solid blue line. Relative actuation speed is the rate of change of *δs′* at the nominal *v*_*CoM*_, assuming no-slip motion.

The snakes could minimize torque by increasing *ξ* and/or *θ*_*m*_, however, we did not observe these waveforms (Fig. 2). Similarly, snakes did not use waveforms which minimized mechanical cost of transport or distance traveled per cycle [21]. We thus hypothesized there were internal factors which “penalized” the low-torque waveforms.

If, for simplicity, we assume the animal moved with no slip regardless of waveform such that the distance traveled per cycle was equal to the wavelength, there were two (not necessarily independent) ways to increase *v*_*CoM*_ : (1) increase the wavelength by decreasing *ξ* and/or *θ*_*m*_ (see example waveshapes in Fig. 7(b)) or (2) increase the speed of wave propagation (given the linear relationship between propagation speed and *v*_*CoM*_, [21]).

In limbed organisms, movement speed is related to the interplay between gait parameters like stride length and frequency and physiological concerns like energetic cost (e.g. [42, 43]). We hypothesized that, because the snake performed an escape response in our experiments, their objective was to maximize speed.

For a desired *v*_*CoM*_, we explored how quickly a “muscle” segment of fixed nominal length relative to total body length would have to shorten as a function of *ξ* and *θ*_*m*_. The shortening speed was a function of both how quickly curvature changed along the body for a given waveshape and the frequency needed to achieve the target *v*_*CoM*_ for that shape’s stride length. We modeled a body segment as a bending beam with length along the spine *δs* and arclength along the inside of the curve *δs*^*′*^ (see diagram Fig. 7(b)). We approximated the amount of shortening the muscles must produce as the difference, Δ*s* = *δs* − *δs*^*′*^, at the point of maximum bending. Using this model we estimated 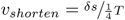, the speed the inside of the bend must change length to go from *δs*^*′*^ = *δs* at the inflection points of the wave to the maximum *δs*^*′*^ occurring at the peaks given undulation period *T* set by the frequency (see Appendix G).

Fig. 7(b) is a heatmap of the increase or decrease in the required shortening speed relative to a nominal shortening speed, *v*_*short,o*_ (which we take to be that occurring at the mean *θ*_*m*_ and *ξ* measured in experiment). The actuator speed increased with *ξ* and *θ*_*m*_ as not only do greater amplitudes and greater wavenumbers require each segment to enact a greater change in body curvature but they also decrease stride length, requiring greater frequency to maintain CoM speed. This is somewhat counter intuitive when considering (non-slip) undulatory gaits as compared to limbed gaits. Increasing stride length on a limbed locomotor requires a greater amount of actuation as, e.g., the hip joint must sweep out a larger angle. Therefore to maintain a CoM speed there is a trade off between increasing/decreasing stride length and decreasing/increasing frequency. In contrast, we see here that actuation speed monotonically decreases as the wavelength increases; if we consider only actuation speed the snakes’ waveforms are under-performing.

This result rationalized why all wave parameters we measured inhabited the center of the wave parameter space (Fig. 2); there is a trade off between external forces and internal actuation constraints.

### B. *C. occipitalis* waveform maximizes movement speed under anatomical constraint

*C. occipitalis* performed an escape response in our trials. Thus, we hypothesized that the specific waveform used by the animals was that which endowed maximum speed across the substrate. We endeavored to include the trade off between decreasing torque and increasing actuation speed needed to maintain *v*_*CoM*_ (now using RFT to estimate the actual stride length given slipping) as *θ*_*m*_ and *ξ* increased. To do so, we used RFT to calculate the joint-level power (the rate of change of the joint angle dotted with the torque experienced by that joint, Appendix F) of each segment over a cycle for each shape in the *θ*_*m*_, *ξ* space. The power-limited velocity, *v*_*pl*_, was the CoM speed for which the peak power generated by any joint over the cycle was equal to a constant, peak available power.

In accord with the tradeoff between internal actuation demands and external torques, *v*_*pl*_ was maximized in the center of the *θ*_*m*_, *ξ* space inhabited by *C. occipitalis’* waveform (Fig. 8(a)). Reflecting the oscillations in *τ*, this metric had maxima near integer values of *ξ*.

**FIG. 8.**
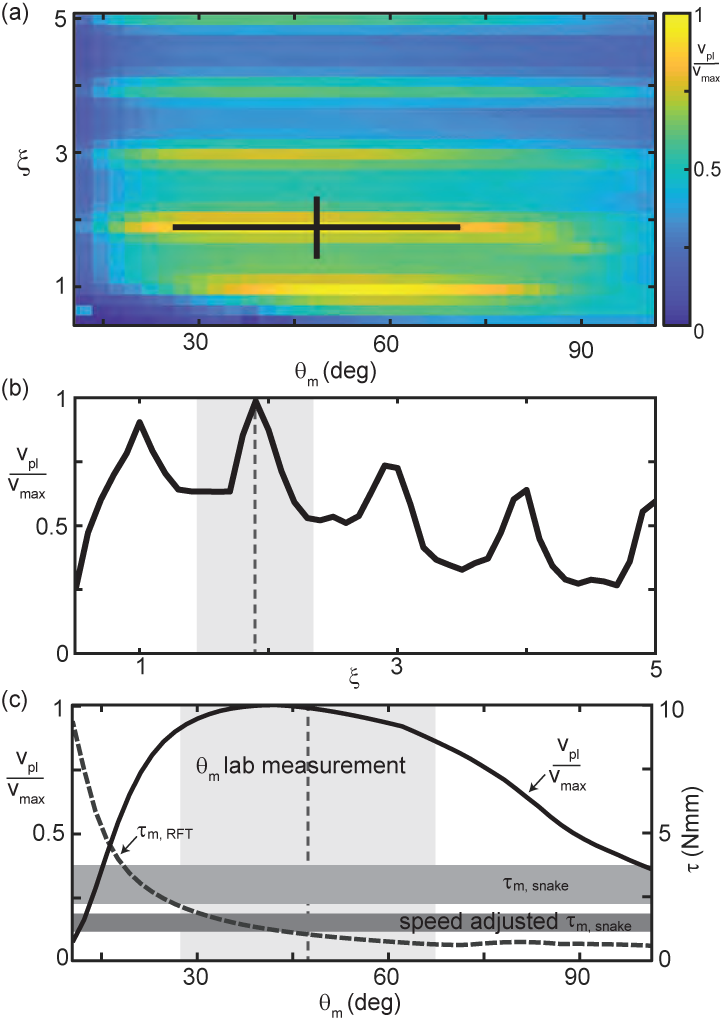
RFT reveals *C. occipitalis’* waveform balances internal and external concerns. (a) Segment-power-limited velocity, *v*_*pl*_, divided by *v*_*max*_, the largest value of *v*_*pl*_, in the *θ*_*m*_, *ξ* space. Black cross is the snake waveform as in (Fig. 2(c)). 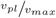 versus *ξ* for *θ*_*m*_ = 48.4, the average experimental value. This is a vertical slice of (a) taken at the average *θ*_*m*_ measured in experiment. The vertical dashed line and gray shaded area are the mean and range of snake *ξ* values. (c) Solid black curve 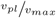 (left vertical axis) and gray dashed curve *τ*_*m,RFT*_ (right vertical axis) versus *θ*_*m*_ for *ξ* = 1.90 (the average snake value) are horizontal slices of (a) and Fig. 7(a), respectively. The vertical dashed line and light gray shaded area are the mean and range of snake *θ*_*m*_. Horizontal gray bars are the estimated muscle torque capabilities of the snakes (Appendix I). The upper bar, *τ*_*m,snake*_ is the estimated maximum muscle torque and the bottom bar, speed adjusted *τ*_*m,snake*_, is the maximum torque reduced to reflect the average muscle shortening speed of the snakes at the average experimentally measured velocity.

Power-limited velocity was maximized at the number of waves (Fig. 8(b)) as well as at the attack angle (Fig. 8(c)) used by *C. occipitalis*. We estimated the peak torque output of *C. occipitalis* muscles using dissection of museum specimens (Appendix I). The attack angle used by the animals was near the point where *τ*_*m,RFT*_ increased above the estimated muscle capability. We report both maximal torque output (Fig. 8(c) upper horizontal bar) as well as output scaled for the average contraction velocity estimated as the body-wall shortening speed (Fig. 8(c) lower horizontal bar). It may be that the individual variation in *θ*_*m*_ reflects differences in peak muscle power capabilities.

### C. Stout snakes must use larger attack angles to succeed

Our granular RFT calculations rationalized the stereotyped waveform used by *C. occipitalis*. The waves of the non-specialists, however, displayed more variation (Fig. 2(c)). Using the insights and RFT calculation we developed in our study of the sand-specialist we endeavored to understand how these differences impacted performance.

A striking difference between the various species was their aspect ratio, *L/w*, the ratio of the total length to the diameter of the body at the widest point. A slender snake like Fig. 1(b,c) will thus have a higher aspect ratio than a stout one, e.g. Fig. 1(a). We previously discovered that *C. occipitalis’* high-aspect-ratio allowed it to move more effectively when swimming subsurface in GM than a low-aspect-ratio sand-swimming lizard [5].

We characterized the performance of the snakes using the unsigned average slip angle, 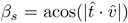 (Fig. 3(a, *inset*), [5]). For each run we calculated slip along the body at each moment in time and took the mean and standard deviation of these values. This variable captures how much the movement of the animal is as if it were moving “in a tube”. If the slip is zero we expect each segment follows in the path of the segment before it whereas when slip is large the body is primarily sliding laterally to the midline.

We found that animals with *β*_*s*_ *>* 30 degrees were those that were ineffective (Fig. 9(a), red crosses). Some snakes with slip close to 30 deg were able to make consistent forward progress (e.g. Fig. 1(e)) while the tracks of those animals with *β*_*s*_ > 30 deg either did not progress at all (e.g. Fig. 1(a,d)), or would have only occasional spurts of forward motion linked by extended periods of undulating in place.

**FIG. 9.**
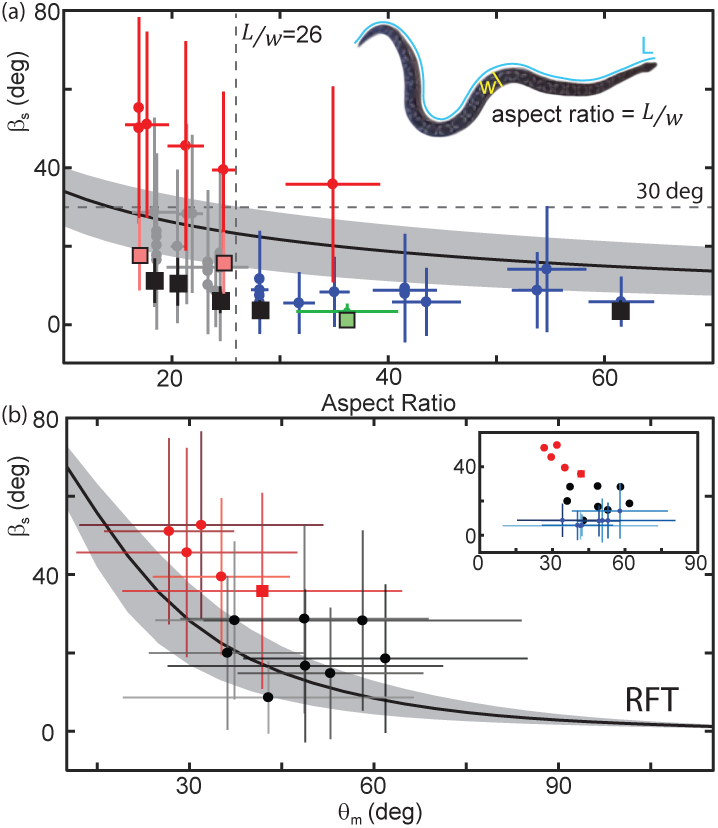
Performance depends on aspect ratio and waveform. (a) Slip as a function of aspect ratio, *L/w* as illustrated on the snapshot of a snake (*A. dumerili, L/w* = 23.2) where L is the total length of the snake (blue curve) and w is the width at the widest point (yellow line). Circle markers and vertical lines indicate the mean and std. of each trial. Horizontal bars are the range of aspect ratios measured from video stills by two different researchers. Black are successful trials, red are failures [21], green is *C. occipitalis*. Each animal tested had a unique aspect ratio; for the non-sand-specialists, multiple data points at the same *L/w* indicate multiple trials for the same animal. N=22, n=38. Only mean shown for *C. occipitalis* (N=9, n=30).Black curve is RFT prediction of slip for the average snake waveform and mass [21] using a scale friction of *µ* = 0.15. Gray area indicates predictions for *µ* = 0.1 (lower slip) and 0.2 (higher slip) (scale friction estimated using [33, 44], Appendix F). Square markers indicate RFT prediction of slip for that animal using a scale friction of *µ* = 0.15 with vertical lines showing range from *µ* = 0.1 0.2. For *C. occipitalis µ* = 0.1 [5] and min/max=0.05/0.15. All RFT predictions used force relations for 300 *µ*m glass beads assuming intrusion depth of 8 mm. (b) Slip versus *θ*_*m*_ for animals with ^*L*^*/w* < 26. Mean and standard deviation of all measurements in a trial. Square marker is the snake *L/w >* 26 which failed. If an animal performed more than one trial we combined the measurements of all trials. Successful trials in black, failures in red. Black curve is the RFT prediction using the average waveform and anatomy values calculated using all species studied, *ξ* = 2.5, mass=0.63 kg, L=89 cm, and an estimated scale friction of *µ* = 0.15. Gray band indicates predictions for friction *µ* = 0.1 and 0.2. (*inset*) Black and red circles are the averages from the main plot. Blue markers are successful trials *L/w >* 26.

We observed a transition in the slip versus aspect ratio plot at an aspect ratio of about 26 (Fig. 9(a), vertical dashed line). At larger *L/w* the snakes were, with the exclusion of one exception, moving effectively with an average *β*_*s*_ = 10.3 ± 3.8 deg (Fig. 9(a), blue crosses). Below *L/w* = 26 performance was highly variable. Four of the five species which failed to progress across the level sand had *L/w* < 26 (Fig. 9(a), red crosses). The other snakes of *L/w* < 26 moved with higher slip on average than their high-aspect ratio counterparts, *β*_*s*_ = 20.3 ± 5.8 deg (Fig. 9(a), gray crosses). This trend was recapitulated in RFT calculation of slip as a function of aspect ratio for a snake of fixed waveform and mass (Fig. 9(a), [21]).

RFT predicted that as attack angle decreased, the slip of a snake with fixed body morphology and wavenumber would increase (Fig. 9(b), black line). We examined slip as *θ*_*m*_ changed for those snakes with an aspect ratio less than 26 (and the snake that failed with *L/w*=30). Those which failed to progress (Fig. 9(b) red crosses) were at lower attack angles than those which succeeded (Fig. 9(b), black crosses)(median and std. of all values *θ*_*m*_(*β*_*s*_ *≥* 30) = 33.2 ± 21.3 N=6, n=4311 measurements, *θ*_*m*_(*β*_*s*_ < 30, *L/w* < 26) = 48.8 ± 24.5 N=12, n=9017, *p* << 0.01). Those snakes which had *L/w* of greater than 26 generally used higher attack angles as well (Fig. 9(b, inset) blue crosses).

RFT was reasonably accurate in predicting the performance of snakes with *β*_*s*_ < 30 deg, given the average *θ*_*m*_, *ξ*, length, and mass of the individual (Fig. 9(a), black and green squares). However, RFT underestimated the slip of those snakes which failed (Fig. 9(a), red squares).

We observed that the snakes created permanent disturbances in the surface of the GM. While those which moved with low slip created observable piles of sand at the posterior-facing side/s of the body (Fig. 1(b,c)), those which failed appeared to dig themselves into a channel, primarily depositing GM laterally to the long-axis of the body (Fig. 1(a)).

## V. MATERIAL REMODELING REGULATES LOCOMOTION

The waveform and performance of the snakes was variable, and this variability was reflected in the tracks left by the animals. As the material did not re-flow around the body after being disturbed, we hypothesized that the manner in which different waveforms remodeled the substrate was important in determining performance. We systematically explored the impact of waveform on performance using a robophysical model, a 10 joint robot on the surface of poppy seeds (Fig. 10).

**FIG. 10.**
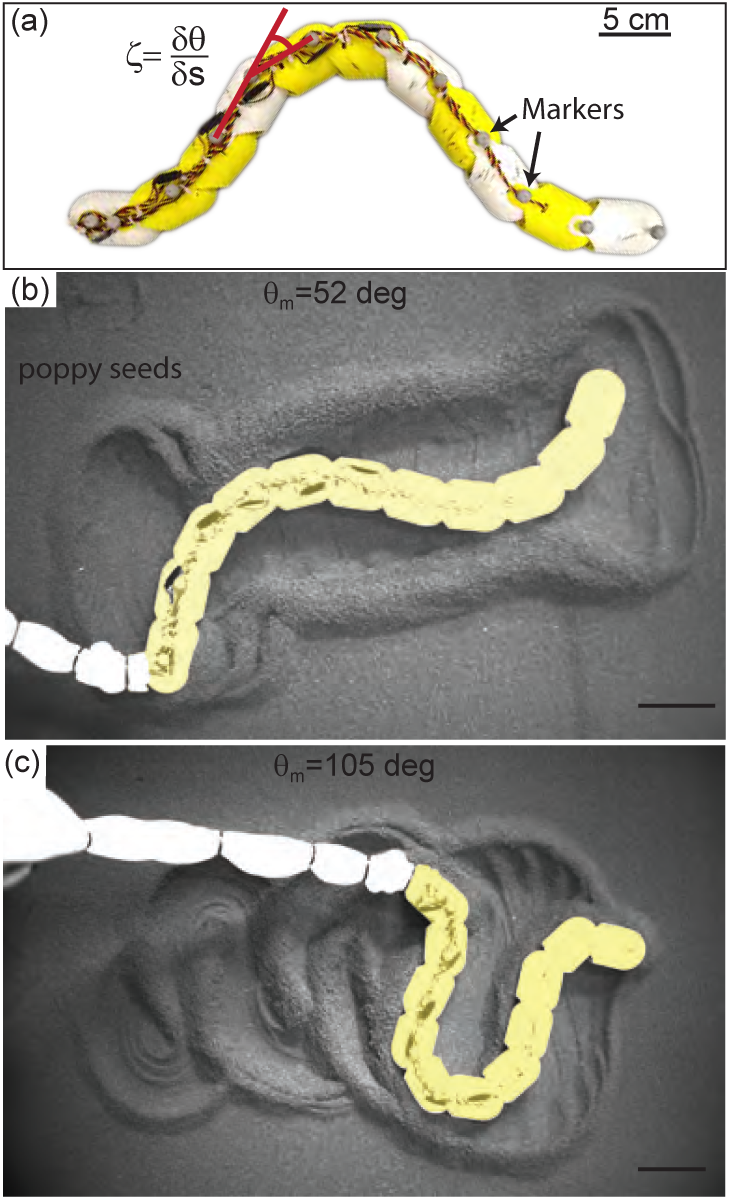
Systematic exploration of memory effects using a robophysical model. (a) 10 joint, 11 segment robot. The shape was a serpenoid curve, 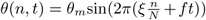 for joint *n* out of *N* total joints, controlled by commanding the servo motor positions, *ζ*, to vary sinusoidally in time with a fixed amplitude. Each motor was offset from its anterior neighbor by a constant phase set by *ξ*. Length=72 cm and *L/w* = 11.3. (b,c) Stills of the robot taken after it has completed 2.75 undulation cycles. *θ*_*m*_ = 52 deg in (b), *θ*_*m*_ = 105 deg in (c), and *ξ* = 1 in both cases. Robot is artificially colored yellow to distinguish the body from the tail cord. Note the large appearance of the tail cord in **c** is due to perspective as the researcher was standing and loosely holding the cord above the substrate.

### A. Robophysical model elucidates connection between waveform and performance

We commanded the motors at each joint of the robot to execute a serpenoid curve (*θ*(*s, t*) = *θ*_*m*_sin(2*π*^*ξ/L*^ (*s* + *v*_*seg*_*t*)) [2]). We varied *ξ* and *θ*_*m*_ and recorded kinematics as the robot performed three undulation cycles on the surface of poppy seeds (Appendix H). The number of joints on the robot was limited by motor strength; as a result the robot could only achieve *ξ* between 1 and 1.4 as lower *ξ* required too much torque and higher were not well-resolved by the number of joints. The power per unit volume of the motors also limited the aspect ratio; *L/w* = 11.3 is lower than that of the stoutest snakes tested.

We characterized performance by measuring slip as in the snakes (Fig. 11(a)). We also measured the CoM displacement in a single undulation cycle normalized by the total length (BLC, Fig. 11(b)). This variable provided intuition for how effectively the robot was progressing across the substrate.

**FIG. 11.**
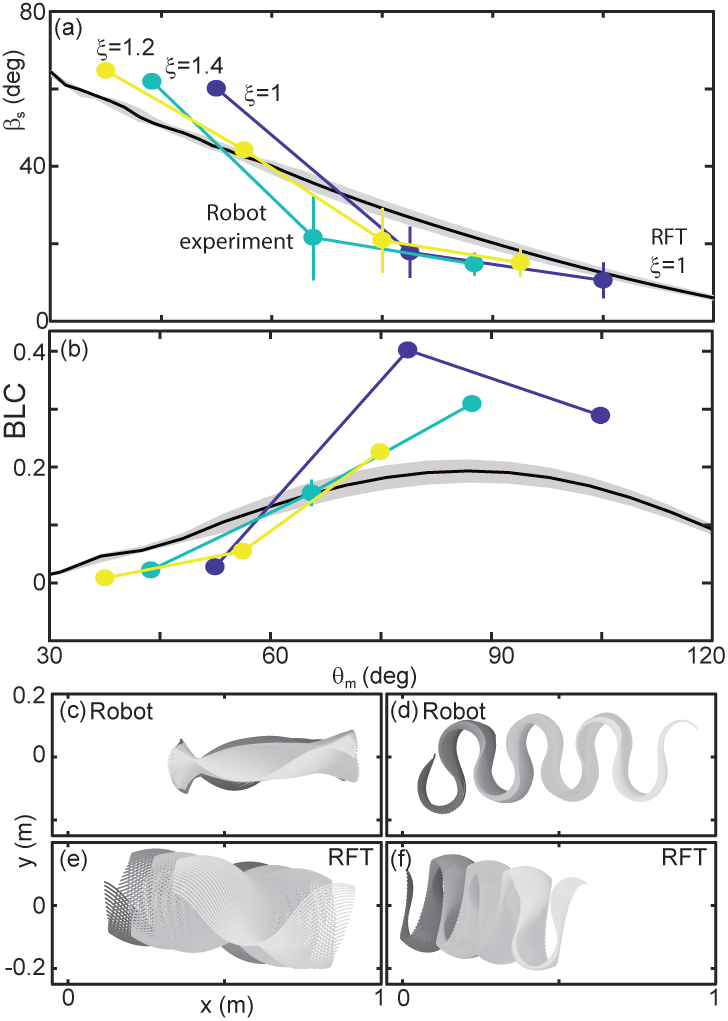
Robot performance is sensitive to the waveform and poorly predicted by RFT. (a) Slip versus *θ*_*m*_. Robot measurements are circles connected by solid lines. *ξ* is indicated by color, yellow is 1.4, teal is 1.2, and blue is 1. Mean and std. of three trials. Where error bars are not visible they are smaller than the marker. RFT prediction is the solid black curve. The gray shaded region indicates uncertainty in our knowledge of the robot’s mass (because of the weight of the tail cord which is held by hand) and friction coefficient. (b) Body lengths traveled in a single undulation cycle (BLC) as a function of attack angle. Colors and lines are consistent with (a). (c,d) Robot and (e,f) RFT predicted kinematics for *ξ* = 1. Color indicates time from the beginning to the end (darker to lighter gray). Both robot and RFT trajectories include three complete undulations. (c,e) *θ*_*m*_ = 52 deg. (d,f) *θ*_*m*_ = 105 deg.

Performance of the robot was a function of the waveform parameters. This was intuitive given our RFT predictions for the snake, which depended on both *ξ* and *θ*_*m*_ (e.g. Fig. 8), and was commensurate with our findings in the biological experiments (Fig. 9(b)). Over the range of *ξ* available to the robot performance was more strongly dependent on attack angle than wavenumber, therefore we focused our attention on variations in *θ*_*m*_.

### B. RFT prediction of robot performance is inaccurate

We used granular stress relations measured in poppy seeds using a flat plate of the same plastic used to print the robot body [24] in a resistive force theory calculation predicting the robot performance as a function of *θ*_*m*_ for fixed *ξ* = 1 (Fig. 11(a,b), black curves).

In previous studies, RFT accurately predicted locomotor performance of a number of systems [15] and in our study RFT was accurate for *C. occipitalis* as well as the successful non-specialist snakes. However, as in the high-slip snakes, RFT under-predicted slip at low attack angles (Fig. 11(a)). Despite the substantial predicted slip, RFT indicated these waveforms would still make forward progress (Fig. 11(b,e)). The robot, however, did not displace at the smallest attack angles tested (Fig. 11(b,c)). Conversely, at high *θ*_*m*_ RFT under-predicted displacement of the robot (Fig. 11(b)).

At both low and high attack angles RFT over-predicted the amount of yaw (rotation about the CoM) that would occur (Fig. 11(c:f)). The discrepancy was likely because this calculation assumed the robot was always encountering undisturbed material. This assumption worked for subsurface swimming where the GM behaves like a frictional fluid, constantly re-flowing to fill in the spaces cleared by motion of the body. We observed that the motion of the robot deposited some material lateral to the direction of motion. These piles, when re-encountered, appeared to limit yaw. Material hysteresis, which was not accounted for in our RFT calculation, appeared to play an important role.

### C. Material remodeling changes performance

The material flow was a function of the waveshape. We characterized the overall structure of this flow by measuring *ψ*_*v*_, the angle between the velocity vector of the grains at a given point in time and space and the overall direction of motion of the robot (Fig. 12(a)). We estimated grain velocities using Particle Image Velocimetry (PIV, Fig. 12(a)), Appendix H) and calculated the probability density to measure a given *ψ*_*v*_ over a run for *ξ* = 1 and each of the three *θ*_*m*_ tested (Fig. 12(b)).

**FIG. 12.**
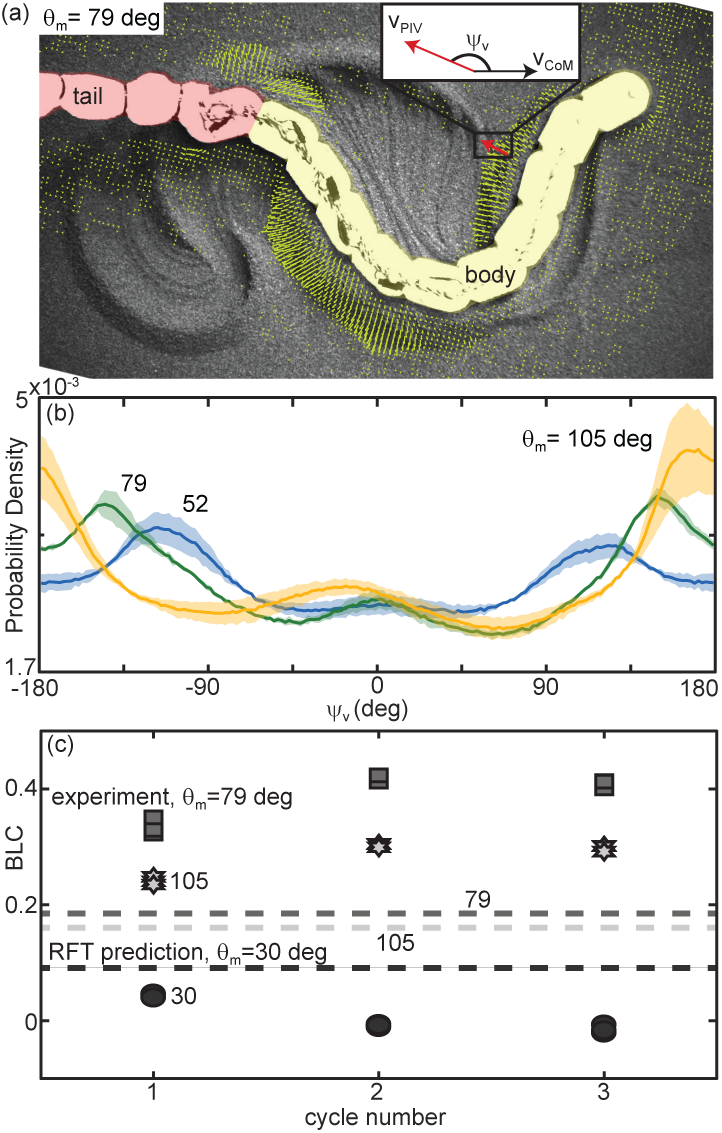
Material remodeling determines kinematics of the robot. (a) Snapshot of the robot moving on poppy seeds with grain velocity vectors estimated using PIV shown in yellow. We measured the grain velocity vector angle, *ψ*_*v*_ as shown. A mask of the body (yellow) and tail (red) was used in all PIV calculations and vectors both too close and too far from the body were not included in calculating *ψ*_*v*_. Vectors shown are representative of those included. We collected three trials per condition and three complete undulation cycles per trial. (*inset*) Diagram of *ψ*_*v*_ for an example estimated grain velocity vector in red. *v*_*CoM*_ is pointing in the average direction of motion of the robot. (b) Probability density of *ψ*_*v*_ for each of the three *θ*_*m*_ tested, *θ*_*m*_ = 105 (yellow), 79 (green), and 52 (blue) as labelled on the plot. Curves were calculated using all *ψ*_*v*_ measured in space and time in a single trial. Solid lines are the average curve and the shaded area indicates the standard deviation across the three trials. (c) Distance traveled by the robot in a single step, normalized by body length, measured at the end of each of three consecutive undulation cycles. *ξ* = 1 for all trials shown. From darkest to lightest gray color: circles are *θ*_*m*_ = 30 deg, square are 70 deg, and stars are 105 deg. Three trials are plotted separately for each step. RFT prediction shown as a horizontal lines with color corresponding to experiment.

At high attack angles significant amounts of the material were deposited by the posterior-facing body segments, similarly to what we observed in the successful snakes (Fig. 10(c), [21]). Consistent with this observation, we measured peaks in the probability density of *ψ*_*v*_ near *±*180 deg, corresponding to grains which are flowing opposite the average direction of motion of the robot (Fig. 12(b), orange curve). When the granular piles moving with the body at later points in time re-enountered these piles the additional stress resisted backward slipping of the robot, increasing displacement above that predicted for a frictional fluid (Fig. 11(b)).

The GM being pushed by low *θ*_*m*_ waveforms had a significant velocity component lateral to the direction of motion; as *θ*_*m*_ decreased, the peaks in the *ψ*_*v*_ probability density shifted toward *±*90 deg (grain flow perpendicular to the average direction of motion, Fig. 12(b), green and blue curves). These piles were thus deposited to the side of the robot and with each undulation more material was added to these “sidewalls” which can be seen in Fig. 10(b). As the GM being pushed by subsequent undulations encountered these pre-existing walls the robot was apparently unable to overcome the inertia of the grains in the previously created piles, instead depositing more material in the walls without changing their location. The result was that the robot swept out a trough in the GM, completely stalling forward progress and sometimes even moving backward as the trunk “rolled” along the trough walls. These failures were similar in appearance to those observed in the living snakes (Fig. 1(a,d), [21]).

In limbed systems, RFT was often useful in predicting the performance of the first gait cycle, before material re-interaction could occur [23, 24]. In the robot, however, RFT was incorrect even for the first cycle (Fig. 12(c)). Enhancement or degradation of the performance was occurring within the first gait cycle as portions of the body contacted GM disturbed by other segments at a previous time.

This provides some intuition as to why *C. occipitalis’* performance was robust to variations in the waveform which would have impacted the robot (Fig. 2); because this animal’s long body and slick scales yield low-slip slithering, the body and/or piles of GM being pushed by the body are continuously encountering new material through time. The animal is effectively moving in a frictional fluid even though the material is not re-flowing as in subsurface movement. This raises an interesting subtlety: the tail is always moving through material disturbed by the head, and placing a successful robot waveform back in its own tracks does not decrease performance [21]. It is apparently the cumulative effect of material deposition coupled with the waveform-dependent pattern of granular reaction forces that conspires to change performance.

Despite the complicated interplay between the robot and the substrate we found the resulting performance was repeatable. For example, for all three attack angles tested at *ξ* = 1, each of the three trials for a given waveform pushed material in a similar way and displaced a similar amount in each of the three undulations (Fig. 12(b,c)). This indicated that, while the material remodeling was complex, it was deterministic.

### D. RFT predicts performance transition

Although RFT did not consistently predict the robot’s slip, it did provide a heuristic for success and give some intuition as to why a locomotor with the morphology of the robot is challenged by GM. The line in the *θ*_*m*_, *ξ* space which RFT predicted to be at *β*_*s*_ ≈ 30 deg separated the successful and unsuccessful robot waveforms (Fig. 13(a)).

**FIG. 13.**
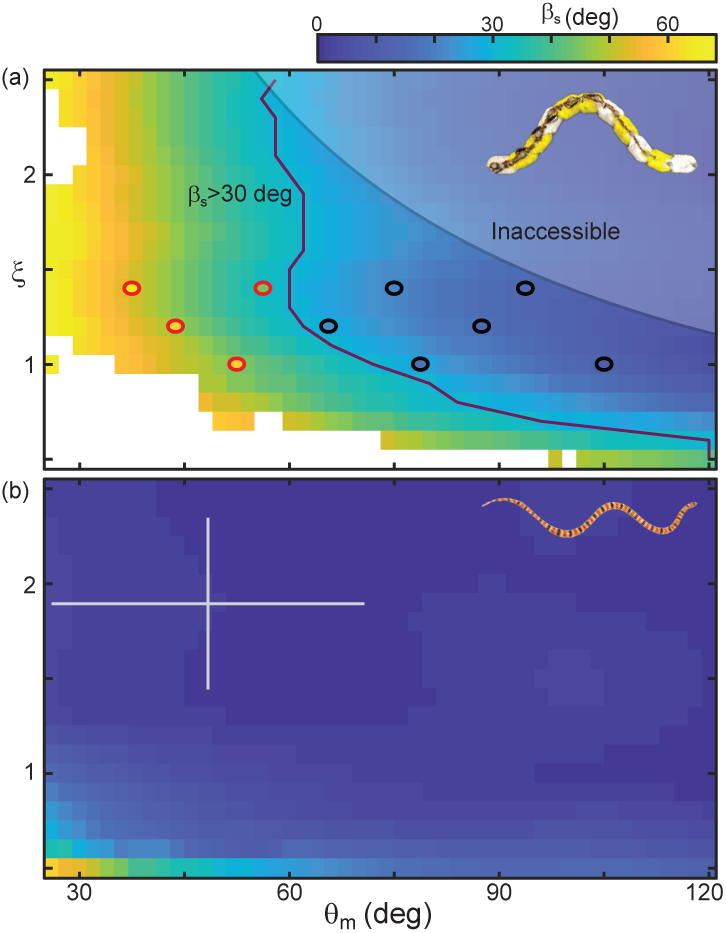
RFT prediction of slip gives a heuristic for performance. (a) Color denotes slip as *θ*_*m*_ and *ξ* vary. RFT is for robot dimensions using poppy seed force relations. Circles indicate the parameter combinations tested on the robot. Waveforms which were successful over three undulations are black, red markers denote unsuccessful parameters (distance per undulation over wavelength *<* 0.35, [21]). Purple line indicates where RFT predicts *β*_*s*_ 30 ≈deg. The lightly shaded region denotes waveforms which the robot cannot perform. (b) Slip calculated for *C. occipitalis* morphology moving on 300 *µ*m glass particles. White cross is animal parameters (mean and range of trial averages) from experiment.

We found there were a limited range of waveforms predicted to both have *β*_*s*_ *≤* 30 deg and be accessible to the robot given its aspect ratio and joint resolution (Fig. 13(a)). In contrast, *C. occipitalis* had a large space of parameters for which slip was small (Fig. 13(b)). This suggests that, in contrast to the robot which was sensitive to waveform, the snake was robust to errors in the motor pattern.

The snake also had the benefit of greater resolution than the robot; the animal has ≈ 120 vertebrae (note that in snakes there is variation in vertebral number between species and between individuals of the same species) while the robot has only 10 joints. This flexibility allowed the snake to operate well away from low-performing shapes (low *ξ* and *θ*_*m*_). Increasing resolution would allow the robot to use more waves on the body, making a greater range of low-slip waveforms available (those in the upper right-hand corner of Fig. 13(a)). However, the robot is limited by the size and strength of the motors. Adding more actuators would increase torque on the central joints, preventing accurate realization of the commanded shapes.

Despite the challenges, locomotion on the surface appears to offer advantages over swimming immersed in material where the fluid-like behavior of the GM allows success of morphologies and motor patterns which fail on the surface [28]. When *C. occipitalis* is on the surface, its escape response is to flee across the GM rather than dive into it [46], suggesting a preference for surface slithering. Drag forces opposing motion are reduced by avoiding the need to push the head through the material as well as decreasing friction on the body which at the surface is due only to animal mass instead of both mass and granular pressure. The drag anisotropy also rises more sharply at the surface, broadening the range of waveforms which can successfully balance thrust and drag forces. Ground contact modulation can further reduce drag forces by lifting segments which are not generating thrust.

## VI. CONCLUSIONS

Undulatory motion on the surface of materials with memory is challenging for both animals and robots. The sensitive connection between performance and waveform is mediated by material remodeling. From study of and comparison between a sand-specialist snake, generalist snakes, and a robophysical model we identified a number of factors which impact undulatory locomotion on our hysteretic material—waveform, aspect ratio, scale friction, and lifting of the wave apexes. The generalist snake and robot experiments demonstrated that possessing all of these is not necessary to motion, but as morphology becomes less optimal (more stout, higher friction) effective management of the substrate remodeling via waveform choice becomes increasingly crucial to progress.

Previous work found that generalist snakes chose self-deformations in response to available environmental forces [6, 27], a strategy which we might expect would be useful on challenging, deformable materials like GM. However, we previously found evidence that *C. occipitalis* does not change its waveform when faced with changes to the surroundings [29] and our robot was executing predetermined waveforms. The ability to sense and respond to induced changes in the substrate is apparently unnecessary given the initial selection of an appropriate pattern of self-deformation.

The interrelationship between morphology, waveform, terrain remodeling, and performance rationalizes why it has been difficult to develop slithering robots which move on terrestrial materials with memory as compared to swimming in fluids or using rigid obstacles. Currently, robots are limited by the power density of the motors to low aspect ratio morphologies with a limited range of available waveforms such that they are in a regime where small changes in waveform can impact progress through the environment. However, despite the intricate nature of the system, we found performance was remarkably stereotyped. This suggests the existence of a predictive model for the relationship between the material changes instigated by body motion and the forces acting between body and substrate. Such a model could both further understanding of the biology and facilitate design of more effective all-terrain robots.

## ACKNOWLEDGMENTS

We thank Kevin and April Young for collecting the *C. occipitalis* and Mark Lowder for his assistance in creating the non-sand-specialist tracking code. This work was supported by NSF PoLS PHY-1205878, PHY-1150760, and CMMI-1361778. ARO W911NF-11-1-0514, U.S. DoD, NDSEG 32 CFR 168a (P.E.S.), and the Simons Southeast Center for Mathematics and Biology.

## Appendix A: Experiments at Zoo Atlanta

Snakes from Zoo Atlanta were tested in an air-fluidized bed of size 2 × 1 m^2^. 200 kg of sand collected in Yuma County, Arizona, USA filled the bed. Between trials a blower was turned on and air flow increased until the GM was in a fluid-like state. The air flow was then decreased until the GM settled in a loose packed state. Air flow was always off during experiments. All experimental procedures were conducted in accordance with the Georgia Institute of Technology IACUC protocol A14058. All procedures involving zoo animals were reviewed and approved by the Zoo Atlanta Scientific Research Committee.

An overhead camera recorded trials at 30 fps and custom MATLAB code identified the snake midlines and fit a 100 point cubic spline at each frame. To measure *ξ* we found the points of zero curvature–corresponding to the inflection points of the waveform. We then measured the arclength between these two points for the first half-wave on the body, multiplied by 2 to get the arclength of one full wave, *λ*_*s*_, and divided the individuals length by the result. We calculated *θ*_*m*_ by first finding *λ*_*s*_ and the wave amplitude in curvature, *κ*_*m*_, to determine the non-dimensional *κ*_*m*_*λ*_*s*_ [5]. *Assuming serpenoid curves*, 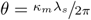. This is a way to estimate *θ*_*m*_ in cases where direction of travel is difficult to determine or the waveform is tortuous.

## Appendix B: *Chionactis occipitalis* experiments

Snakes were collected by Kevin and April Young in the Mojave desert in Yuma, Arizona, USA under scientific collection permits (SP790952, SP625775, SP666119) approved by the Arizona Game and Fish Department and held in the Physiological Research Laboratory at Georgia Tech. Neither the sex nor the age of the animals was determined; gender and age dependent effects were beyond the scope of this study. All experimental procedures were conducted in accordance with the Georgia Institute of Technology IACUC protocols A14066 and A14067.

Glass-oxide particles (Ballotini Impact Beads, Potters Industries Inc.) filled an air-fluidized trackway of area 152 × 53 cm^2^ to a depth of ≈ 4.5 cm. The bottom cavity of the trackway was connected to a blower (Dayton 10 5/8” wheel diameter) which forced air through a layer of Al honeycomb (Plascore, Zeeland, MI USA. PAMG-XR1-8.1-1/8-002-N-5052-B09) and porous plastic (Interstate Specialty Products, Sutton, MA USA. 0.2” thick 50-90 *µ*m pore size) which both supported the sand and acted as a flow distributer.

We fluidized the GM between each trial to erase tracks and reset the substrate to a featureless, loose-packed state; air flow was turned off during trials. The temperature in the track way and snake holding area was measured prior to each trial. Lamps were used to ensure the temperature in both remained at 26 *±* 1 °C, within the active range for *C. occipitalis* [47]. The heat lamps on the track way were turned off during data collection and LED lights were used for illumination.

An overhead high-speed camera (AOS Technologies X-PRI or S-Motion, Baden Daettwil, Switzerland) captured the motion of the snakes at 250 frames-per-second. Each day the individuals to be tested were transported from the housing facility to the lab where we conducted the trials. Snakes which were in the process of shedding were not used. During a trial, the snake was removed from its holding container and placed immediately in the fluidized track way. The snakes tended to be skittish and handling both during trials and in the housing facility was kept to a minimum. The animals would often immediately flee across the surface upon introduction to the track way; otherwise a light tail tap would elicit an escape response. If an individual did not respond to this stimulus they were returned to the holding container. If an individual failed to perform for three trials in a row they would be retired from the days studies. Snakes were tested at most every other day with a maximum of two successful trials collected per day. A run was included if the snake performed at least four complete undulations moving along a straight trajectory at apparently constant speed.

We tracked the black bands on the snake to determine the body midline positions using the method described in [5]. The number of tracked points depended on the number of bands on an individual. To facilitate comparison between snakes of different lengths, a cubic spline was fit to the tracked data such that each snake was evenly divided into 100 measurements from neck to vent.

*θ* was calculated using finite differences to estimate the x and y values for each segments tangent vector. We generated sample serpenoid waveforms of known parameters and used these to determine that, in the presence of noise, the most accurate measurement of the tangent vector was obtained by subtracting the average position of the segment in question and three segments anterior from the average of the segment and the next three posterior segments. This method helped buffer against noise while still providing an accurate measurement. We found the maximum angle on the body at each time step and then averaged over all times in a trial in the to obtain *θ*_*max*_. The average *θ*_*m*_ we measured in experiment was comparable to that measured from photographs we collected of tracks in the desert (*θ*_*m*_ = 45.0 *±* 12.7°, *P* = 0.31 [21]). *ξ* was found using the same algorithm as in the zoo snakes.

## Appendix C: Lifting measurement

An OptiTrack motion capture system (four Prime 17W cameras) captured 3D kinematics of the animals moving in the track way (see [21] for calibration). OptiTrack’s Motive program calculated the x,y, (horizontal plane) and z (vertical distance above the GM surface) coordinates of IR-reflective markers 3 mm in diameter placed at 1.5 cm intervals along the mid-line of six of the *C. occipitalis*, starting between the eyes and continuing to the vent. The markers were small enough we did not observe any difference in the snakes behavior or movement after application.

Space-time plots of *θ* (the horizontal wave on the body which is clearly seen by eye) and z (the vertical wave of lifting) show traveling waves in both. The wave of lifting in z appeared to be double the spatial and temporal frequency [21]. We fit the function *a*_*s*_sin(*ξs*) + *b*_*s*_cos(*ξs*) to each time step in *θ* and z, separately. We included only those values for which R-squared≥ 0.70. The median value of *ξ* for the horizontal wave was 1.6 *±* 0.2 and for the vertical wave 3.4 *±* 0.5. The ratio of vertical to horizontal waves was 2.1, that is, the snake lifted the body segments twice for every one wave passed down the body. Note that *ξ* of the horizontal wave is less than that reported in the text for the animals. This was because in measuring the *ξ* of the animals we divided the total length of the animal by the arclength of the first complete wave. In contrast, here we are fitting the data points which do not represent the entirety of the snakes body as some amount is lost in averaging to calculate *θ* and the posterior points were frequently missing as sand would flow over the tail.

We fit the function *a*_*t*_sin(*ωt*) + *b*_*t*_cos(*ωs*) to the trajectory of each point through time for *θ* and z, separately. We found the wave phase by converting the amplitudes *a*_*t*_ and *b*_*t*_ to polar coordinates and taking the polar angle. A plot of the phase of *θ* compared to that of *z* reveals two linear bands [21], indicating the propagation of the waves is linearly related. The wave of lifting occurred twice as fast as the horizontal wave. Further, we note that the wave of lifting is in turns in phase and *π* out of phase with the horizontal wave. Peaks of the lifting wave align with *θ* = 0 (the highest-curvature portions of the wave).

Given uncertainty in the surface we could not determine whether the animal was in contact with the surface or not. However, we assumed that lifted segments are applying less load to the material and the track depth measurements suggest that the apexes are not in contact.

## Appendix D: Laser line measurements of the free surface

We characterized the tracks created by *C. occipitalis* using a laser-line apparatus. An off-axis camera (Logitech C920 HD Pro Webcam) captured images of the laser line. Both the laser and camera were affixed to a linear bearing and moved using a linear actuator (Firgelli). We used MATLAB to move the actuator and collect an image every 1 mm, verified using a ruler placed in view of the camera. The location of bright pixels in the images was used to reconstruct the surface [21].

## Appendix E: Granular drag measurements

The same glass particles used in the *C. occipitalis* experiments filled an air-fluidized bed of volume 32.5×28×15 cm^3^. The bottom cavity of the bed was fluidized by two shop vacs controlled by a proportional relay. The fluidization mechanism was the same as in the snake trials, and again the material was fluidized before each trial and the air was turned off and all grain movement ceased before a trial began. An aluminum rod attached the Al plate to a force transducer (ATI nano43) accurate to 2 mN. This apparatus was located on the end effector of a six degree-of-freedom robot arm (Denso VS087A2-AV6-NNN-NNN) which controlled intruder movement. The robot arm first rotated the plate to *β*_*s*_, then submerged it to depth *z*, next dragged the plate parallel to the surface at constant speed for 20 cm, and lastly stopped and extracted it. Wherever a mean and standard deviation is given this was taken over data taken at 10 to 20 cm drag distance, the point at which the force had reached steady state and the robot arm was moving at the commanded velocity.

## Appendix F: RFT formulation

We observed that as a body segment pushed laterally against the sand it built a pile similarly to the pile that evolved during drag of the plate. The rostral-facing side of the body, opposite the pile, appeared to be in contact with very little material. Therefore, we chose to model the snake body as a flat plate, representing the caudal-facing side of the body, with an added term to account for drag on the ventral surface. This is not be an unreasonable picture of this species, as the ventral surface is known to be flat or even slightly concave [31]. As the snake held the head slightly raised from the surface we did not include a term accounting for the drag acting against the head as it is pushed through the material.

Previous research [33] found that the ventral scutes are anisotropic such that gliding directly forward produces less drag than sliding laterally and both forward and lateral motion results in less frictional drag than moving directly backwards. The coefficient of static friction of C. occipitalis scales is low (0.109*±*0.016 ventral forward and 0.137*±*0.018 backward [5]). The lateral coefficient of friction for *C. occipitalis* scales is not known, however, given [33] results we assumed that it was bounded between 0.10 and 0.14 such that the difference between frictional forces acting tangential and normal to segments was negligible compared to the larger plate-drag forces. Therefore, we chose not to include the frictional anisotropy in our RFT calculation. We previously found the coefficient of static friction between Aluminum and the glass particles is approximately 0.2 [32]. Therefore, in predicting the performance of *C. occipitalis* we scaled the *σ*_*t*_ function measured using the Aluminum plate by ½.

In modeling the various snake species we used frictions 0.1, 0.15, and 0.2. This represents a reasonable range of the friction coefficient between forward-sliding ventral scutes and the substrate during lateral undulation [33, 44], although we note that higher coefficients have been measured on rough material and can be actively modulated by the animal [48]

We modeled ventral drag on each segment as acting opposite the velocity with magnitude *µmg/n*_*segs*_ (*µ* = 0.1 was the coefficient of static friction for the snake’s ventral scutes on the glass particles [5], *m* was the average snake mass, *g* = 9.81 m s^2^ was the gravitational constant, and *n*_*segs*_ the number of segments used in the calculation).

Given the low-friction scales of *C. occipitalis*, including lifting in the calculations did not substantially improve predicted performance [21]. We speculate that these segments may be lifted as a side-effect of the muscle activation responsible for generating the horizontal waveform, that is, the morphology of the trunk means that vertical lifting is a consequence of generating the curvature in the horizontal waves. It may also be that the lifting is intentional and a buffer against deleterious motor program mistakes or changes in the environment, or the small benefit of lifting these segments is greater than the energetic cost. We did not re-distribute ventral drag forces to account for lifting.

We approximated the kinematics of the snake using 100 segments at 70 points in time divided evenly over one full undulation cycle. These values are well past those needed for convergence of the calculation. We calculated each segment’s orientation and velocity using 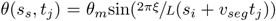 for each combination of parameters *θ*_*m*_ and *ξ*. Summing over the body yielded the total force acting on the CoM in *x* and *y* and torque about the CoM. MATLABs lsqnonlin function was used to find the *x* and *y* components of *v*_*CoM*_ and the angular velocity about the CoM which resulted in zero net stress.

We calculated torque acting on the CoM using 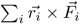 where 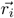 is the location of each segment with respect to the CoM and 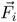 are the RFT calculated forces acting on that segment. We calculated torque acting at each joint as in [6]. For joint number *n* the internal torque is 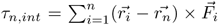. To predict *τ*_*m,RFT*_ in a cycle we find this value for each of the segments at all times. *τ*_*m,RFT*_ is the maximum value in this 100 × 70 matrix. Joint-level power for each joint at each time was calculated using the rate of change of the joint angle, 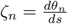,

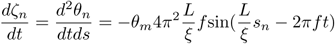

and power was thus

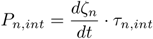

## Appendix G: Bending beam model for snake segment motion

We kinematically model a snake body segment as a beam with midline length *δs* and length of the inside of the curve *δs′*. For a given body radius, *r*, and radius of curvature, 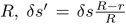. The speed the inside of the segment must change length from the unbent length of *δs* to *δs′* is 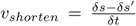. We can write *δs - δs′* as 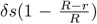. Using the relation *R* = *κ*^*-*1^ we get *v*_*shorten*_ = *δsrκ*. Next, using 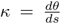 for a serpenoid curve 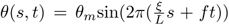, we find the maximum curvature in terms of attack angle 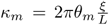. Thus the maximum length change 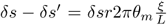. The time the segment has to undergo this change is a quarter of a period, so *δt* = (4*f*)^*-*1^. Thus the shortening speed of the inside of a segment is

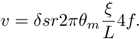

Lastly, for the nominal values *f*_*o*_, *ξ*_*o*_, *θ*_*m,o*_ we get the ratio 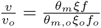 and the relative shortening speed is

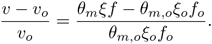

For a predetermined *v*_*CoM*_, MATLAB’s fsolve is used to find the undulation frequency for a given *θ*_*m*_ and *ξ* which results in no-slip motion at that value of *v*_*CoM*_.

## Appendix H: Robophysical Experiments

Robot experiments were performed in a 1.2×1.8 m^2^ bed filled to a depth of 10 cm with poppy seeds, manually smoothed between trials. For the kinematics trials four OptiTrack cameras (Flex 13, Natural Point) tracked infrared reflective markers on the robot. PIVlab (v1.41) [50, 51] was used to estimate grain velocity measurements from video taken using an overhead camera (AOS S-Motion, Baden Daettwil, Switzerland) captured video at 120 fps.

Robot was constructed using ten Dynamixel AX-12A servo actuators (Robotis) with a stall torque of 15.3 kg*·*cm. Each motor was housed in a cylindrical, 3D printed casing. Joint positions were commanded using a Lynxmotion SSC-32 servo controller.

## Appendix I: Dissection and torque estimation

Four adult Chionactis (SVL 33.0 *±* 3.7 cm, mass 16.4 *±* 3.4 g) which had preserved in formalin and stored in ethanol were scalened and the body (from the posterior margin of the quadrates to the cloaca) was cut into 10 equal length segments, which were then weighed intact. Segments were then eviscerated and the epaxial muscle mass (consisting of the largest muscles, the m. multifidus, m. semispinalis-spinalis, m.longissimus dorsi, and m. iliocostalis [52]) was removed, and the viscera, muscle, and remaining body tissue were weighed. All tissues were kept moist in 70% ethanol and dabbed dry before weighing. Snakes were an average of 22.1 *±* 3.8% muscle by mass, but this proportion varied regionally due to uneven total segment masses and viscera masses; absolute muscle mass was highest at midbody and decreased anteriorly and posteriorly, though the highest muscle mass was only 27.0 *±* 3.8% higher than the average.

Because of postmortem and preservation distortion of body shape (facilitated by the mobile ribs of snakes), average body radius was computed from SVL and mass by treating the snake as a uniform cylinder with a tissue density of 1.05 g cm^3^ (typical for vertebrate tissues). As the muscular lever arms for lateral flexion are unknown for any snake, we estimated the maximum lever arm as ^1^*/*2 the radius of the body; while some epaxial muscles may have larger lever arms (m. iliocostalis), others likely have much lower lever arms (m. multifidus and semispinalis-spinalis). Similarly, muscular PSCA for the entire epaxial muscle group was computed as a cylinder based on SVL and total muscle mass; while snake epaxial musculature is highly complex, none of the muscles show strong pennation. Peak isometric muscle force was estimated based on the standard 30 N cm^2^ value seen in most vertebrate muscles, and divided by two to account for unilateral activation [52]. Although the maximal shortening speed and shape of the force-velocity relationship is unknown in snakes, we assumed that during lateral undulation, snakes would be operating near their peak isotonic power, and thus with a force of half the peak isometric muscle force; as activation/deactivation kinetics and length tension properties are also unknown in snakes, we did not attempt to account for these. Peak torque was computed as this force divided by estimated lever arms. Although many crucial properties are unknown, this value represents a charitably high estimate of peak torque; this value would be depressed by steeper force-velocity curves, departure from the plateau of the length-tension curve, incomplete activation during the work loop, or lower muscle lever arms, while the higher muscle mass at midbody and slightly larger vertebrae would increase the peak torque.

## SUPPLEMENTARY INFORMATION

### 1. *C. occipitalis* husbandry

Animals were housed individually in ten-gallon glass aquariums filled with dry sand to a depth of approximately 10cm and were provided a water dish and a moist hide made of a small plastic container filled with damp moss. A heat pad was adhered to one end of the tank to provide a temperature gradient. The housing room was on a 12 h:12 h light:dark schedule. Snakes were given 6 crickets coated in a supplemental calcium powder three times a week and allowed to eat ad libitum.

### 2. PCA

Inspired by [1] we used principal component analysis (PCA) to search for a low-dimensional representation of the waveform. There were two dominant PCs which were well-fit by sinusoids with phase difference *π/*2 (Fig. S1(b)). These two vectors accounted for ≈ 91% of the variance, such that the waveform was well-approximated by

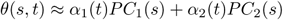

A plot of *α*_1_ versus *α*_2_ revealed the coefficients trace out a circle in time (Fig. S1(c)) which, when combined with the sinusoidal PCs, corresponds to a traveling sinusoidal tangent angle along the body. Such a wave is called a serpenoid curve, 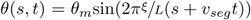 [2]. The amplitude *θ*_*m*_ is the maximum value of *θ, ξ* is the number of waves on the body, and *v*_*seg*_ is the wave propagation speed of the segments (the speed of travel if one were riding on the body).

### 3. Wave apex lifting measurements

We calibrated the origin of the Optitrack coordinate system using the provided calibration square. We placed the square so that the top surface was level with the free surface of the GM and used a level to ensure the square was horizontal. The error in placing the square was difficult to determine but we estimate it was within 3° in any direction. Because the calibration was imperfect we rotated the z-coordinates of the snake by fitting a line to the average position of the snake in y and z at each moment in time, finding the angle between y and z given the slope of this line, and using a rotation matrix to rotate the z coordinates.

### 4. RFT calculations

While shape effects on drag at the surface are not well-known, during subsurface drag the horizontal forces on an object depend primarily on cross-sectional area with very little dependence on the shape of the object, much less than in fluids [3]. While there is a shape-dependence on forces in the vertical direction [4], we assumed that vertical forces acting on the snake were balanced and did not affect the horizontal forces determining performance.

#### a. RFT prediction of mCoT and BLC

When swimming subsurface *C. occipitalis* waveform minimized mechanical cost of transport (mCoT), calculated as the mechanical power needed to complete one cycle divided by the product of CoM velocity and mass [5]. We used surface RFT to calculate mCoT as *θ*_*m*_ and *ξ* varied (Fig. S7(a)). mCoT increased when *θ*_*m*_ or *ξ* were small as to achieve zero net stress these shapes required segments to be at larger values of *β*_*s*_ therefore yielding more material and requiring greater power. Shapes with high *θ*_*m*_ also increased mCoT as the body spends more time going “up and down” rather than making forward progress, reducing *v*_*CoM*_. There was a large basin of minimal mCoT at intermediate values. The white cross indicates the mean and range for all trials. While was a large range of *ξ* which minimized mCoT the snake used only a limited subset of these. The animal’s *θ*_*m*_ was also somewhat higher than that which would yield minimal mCoT.

We predicted distance traveled per undulation cycle (dimensionless quantity body-lengths per cycle, BLC). Given the linear relationship between *v*_*seg*_ and *v*_*CoM*_, a waveform which maximized BLC would afford the greatest increase in *v*_*CoM*_ per increase of *v*_*seg*_. BLC decreased as the number of waves on the body increased (Fig. S7(b)). The body length was fixed, so adding waves on the body shortened “stride length” Changing *ξ* did not change mCoT as drastically because decreasing the arc length of a single wave (by increasing *ξ*) decreased both power and *v*_*CoM*_. The animals appeared to be using a waveform which increased BLC in comparison to the subsurface shape (crosses in (Fig. S7(b)), however, BLC was maximized at a lower *ξ* than used by the animal.

We did not include in our calculation the energy required to overcome internal resistance from viscous and/or elastic stresses of the snake body. [6] reported the internal resistance of the sandfish was an order of magnitude less than that of the external resistance of the GM during movement subsurface. The GM stresses at the surface were less than subsurface by about an order of magnitude and the snake cross-sectional area is about half that of the sandfish (snake radius ≈ 5 mm versus sandfish ≈ 7.5 mm, assuming circular cross section). The stiffness and damping of an elastic rod is proportional to cross-sectional area. Thus, while the difference between internal and external stresses is likely not as large as during subsurface motion, we chose to ignore the internal stresses. Future study could combine RFT with a model for the internal properties of the animal’s body following, e.g., [7].

#### b. RFT for changing aspect ratio and mass

We used the average mass m=0.76 kg, length L=0.89 m, *θ*_*m*_ = 48, and *ξ* = 2.38 of all non-sand-specialist snakes with slip *<* 30 deg. We assumed constant body segment length (i.e. all snakes divided into same number of segments), constant body density, *ρ* = 1000 kgm^*-*3^, and a circular cross section. The width and length of a snake can then be calculated as

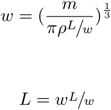

The force on each segment is assumed to scale linearly with area, so the granular force relations are both scaled by a constant

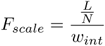

where N=100 is the number of body segments and *w*_*int*_ is the width of the Aluminum plate used in drag experiments. We assumed that the intrusion depth of the snake body was equal to that of the plate, *z* = 8 mm.

---

**TABLE S1.** Anatomical information for the non-sand-specialist species.

| Species | subfamily | family | length (cm) | max width (cm) | mass (g) |
| --- | --- | --- | --- | --- | --- |
| *Acranthophis dumerili* |  | Boidae | 183 | 7.9 | 5620 |
| *Agkistrodon bilineatus* | Crotalinae | Viperidae | 75.4 | 3.3 | 306.7 |
| *Agkistrodon contortrix* | Crotalinae | Viperidae | 80 | 3.1 | 359.5 |
| *Agkistrodon piscivorous* (1) | Crotalinae | Viperidae | 92 | 4.5 | 569.5 |
| *Agkistrodon piscivorous* (2) | Crotalinae | Viperidae |  |  | 874.5 |
| *Agkistrodon piscivorous* (3) | Crotalinae | Viperidae | 57 | 2.7 | 161 |
| *Aspidites ramsayi* |  | Pythonidae | 1.3 | 3.4 | 749 |
| *Bothriechis schlegelii* | Crotalinae | Viperidae | 66.7 | 1.7 | 104 |
| *Crotalus lepidus* | Crotalinae | Viperidae | 43 | 2.2 | 44 |
| *Crotalus molossus* | Crotalinae | Viperidae | 82.1 | 4 | 370.7 |
| *Crotalus tigris* | Crotalinae | Viperidae | 71.1 | 3.1 | 331.6 |
| *Crotalus willardi* | Crotalinae | Viperidae | 47 | 3 | 134.5 |
| *Dasypeltis scaraba* |  | Colubridae | 71 | 1.1 | 51 |
| *Drymarchon couperi* |  | Colubridae | 162 | 4.2 | 1018 |
| *Epicrates subflavus* |  | Boidae | 153 | 3 | 738 |
| *Lampropeltis getula getula* | Colubrinae | Colubridae | 121 | 3 | 680 |
| *Lichanura trivirgata* |  | Boidae | 71 | 2.6 | 243.5 |
| *Loxocemus bicolor* |  | Loxocemidae | 111 | 3.3 | 607.5 |
| *Nerodia sipedon sipedon* | Natricinae | Colubridae | 79 | 3.5 | 453.5 |
| *Senticolus triaspis* | Colubrinae | Colubridae | 101 | 1.8 | 198 |
| *Sistrurus catenatus* | Crotalinae | Viperidae | 52 | 2.8 | 174.5 |
| *Sistrurus miliaris* | Crotalinae | Viperidae | 47 | 2.8 | 146 |

**TABLE S2.** Anatomical information for the individual sand-specialist snakes. The species *Chionactis occipitalis* is in the family Colubridae.

| Individual | length (cm) | max width (cm) | mass (g) |
| --- | --- | --- | --- |
| 120 | 36.4 | 1.1 | 20 |
| 122 | 39.2 | 1.0 | 21 |
| 123 | 40.1 | 1.1 | 20 |
| 124 | 38.1 | 1.1 | 20 |
| 125 | 36.6 | 1.0 | 18 |
| 128 | 38 | 1.0 | 16 |
| 129 | 39.3 | 1.2 | 24 |
| 130 | 37.1 | 1.0 | 20 |
| 132 | 37.1 | 1.0 | 18 |

**FIG. S1.**
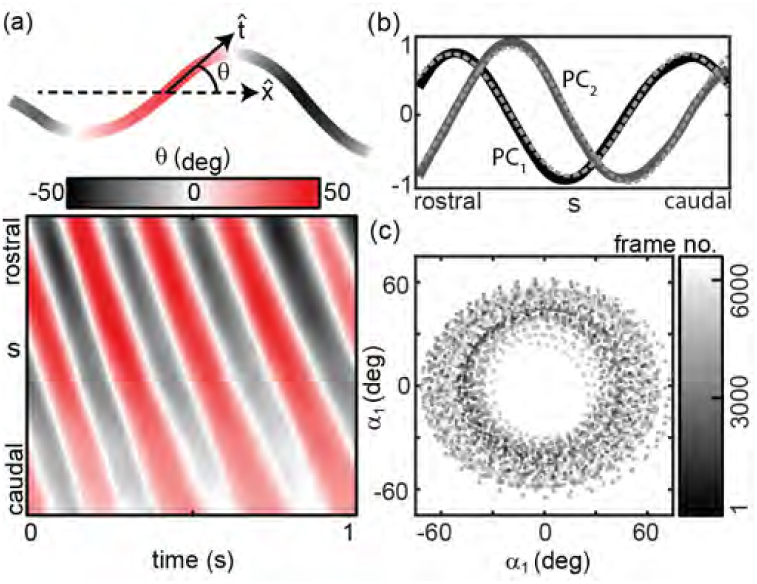
Principal component analysis (PCA) of all sand snake shapes. (a, *top*) Tangent angle, *θ* is the angle between 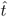 and the average direction of motion of the animal,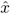. (*bottom*) Example space-time plot of *θ*, corresponding to the *bottom* trial in c. The diagonal bands indicate a posteriorly traveling wave propagated at constant speed. (b) The two dominant PCs representing *θ*. The solid lines are the experimental results, the dashed lines are sine fits to *A*_*i*_sin(*ξ*_*i*_*s* + *ϕ*_*i*_). Coefficients with 95% confidence bounds were *A*_1_ = 0.83 (0.82, 0.84), *A*_2_ = 0.92(0.90, 0.94), *ξ*_1_ = 1.44(1.43, 1.45), *ξ*_2_ = 1.40(1.39, 1.41), *ϕ*_1_ = 0.05*π*(0.04*π*, 0.06*π*), *ϕ*_2_ = −0.41*π*(0.42*π*, 0.39*π*). *ϕ*_1_ −*ϕ*_2_ = 0.46*π*. Both vectors were divided by the same maximum value such that the maximum amplitude was one. (c) The amplitudes *α*_1_ and *α*_2_ associated with the PCs in e. Color indicates the frame number for all trials (N=9, n=30) combined. The trajectories move counterclockwise around the circular structure, indicating a traveling wave. The radius of this circle is the amplitude of the wave of *θ*.

**FIG. S2.**
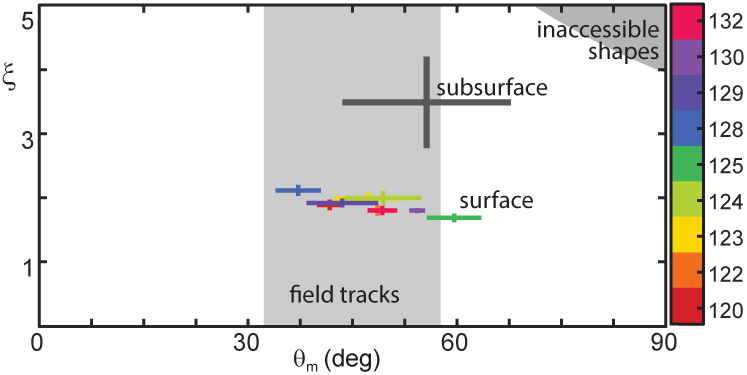
*C. occipitalis* kinematics compared to field measurements. (a) (Plot as in main text with field measurements added). The light gray rectangle illustrates the range of *θ*_*m*_ measured from photographs taken of *C. occipitalis* tracks found in the desert (17 measurements). Experimental measurements plotted in the serpenoid curve parameter space {P*θ*_*m*_, *ξ*}. Color indicates animal number. Markers are the mean and range of each individual taken over all trials. N=9 individuals, 30 trials. The gray cross is the waveform used by *C. occipitalis* when moving subsurface. The surface waveform was distinct from that used by this species when moving buried within the GM [5]). While the angular amplitude was not significantly different (subsurface *θ*_*m*_ = 54.7 ± 11.9° compared to the surface value *θ*_*m*_ = 48.3 ± 7.0° for all snakes combined, 2-tailed t-test *P* = 0.04 mean of all trials and std of mean of all trials), the value of *ξ* was less for surface swimming (*ξ* = 3.53 ± 0.85 subsurface versus *ξ* = 1.90 ± 0.14 at the surface, P*<* 0.001). The gray region in the upper right corner are waves which are inaccessible given the flexibility of the snake [5].

**FIG. S3.**
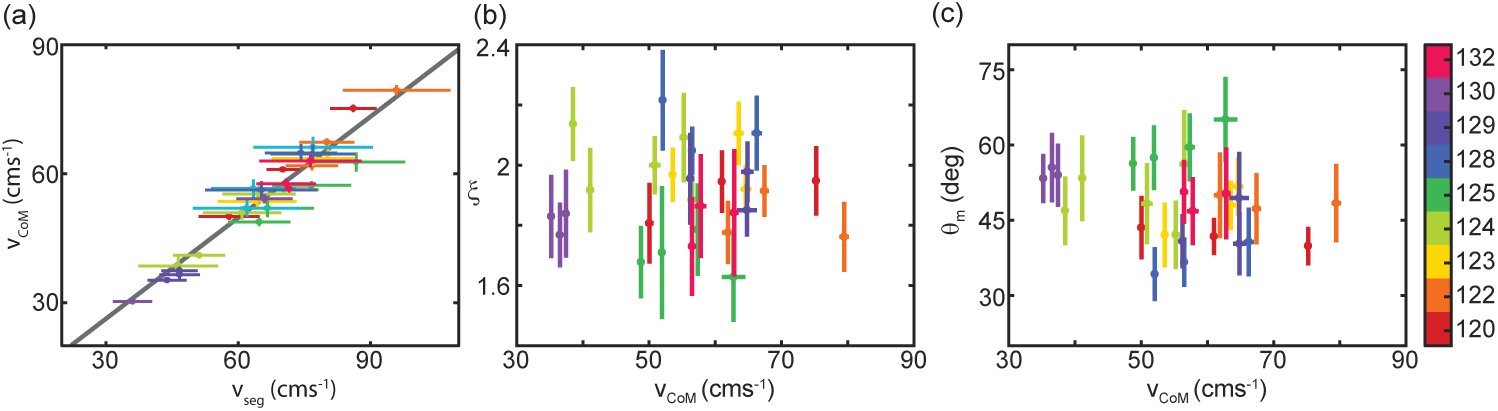
(a) Center-of-mass velocity, *v*_*CoM*_, versus segment velocity (as if one were riding on the body), *v*_*seg*_. Lines are mean and std. of each measurement. Each data point is an individual trial, individual number as shown in colorbar to the far right. Colors are consistent with those in main text Fig. 2. Gray line is the linear fit (mean and 95% confidence intervals of slope=0.81 (0.74, 0.89), y-intercept=0.67 (−4.63 5.94), R-squared=0.94, RMSE=2.88) N=10 individuals, 32 trials. N=10 individuals, 32 trials. (b) Number of waves on the body, *ξ*, versus *v*_*CoM*_. Mean and std. for each trial, colored according to individual. (c) Attack angle, *θ*_*m*_, versus *v*_*CoM*_. Mean and std. for each trial, colored according to individual.

**FIG. S4.**
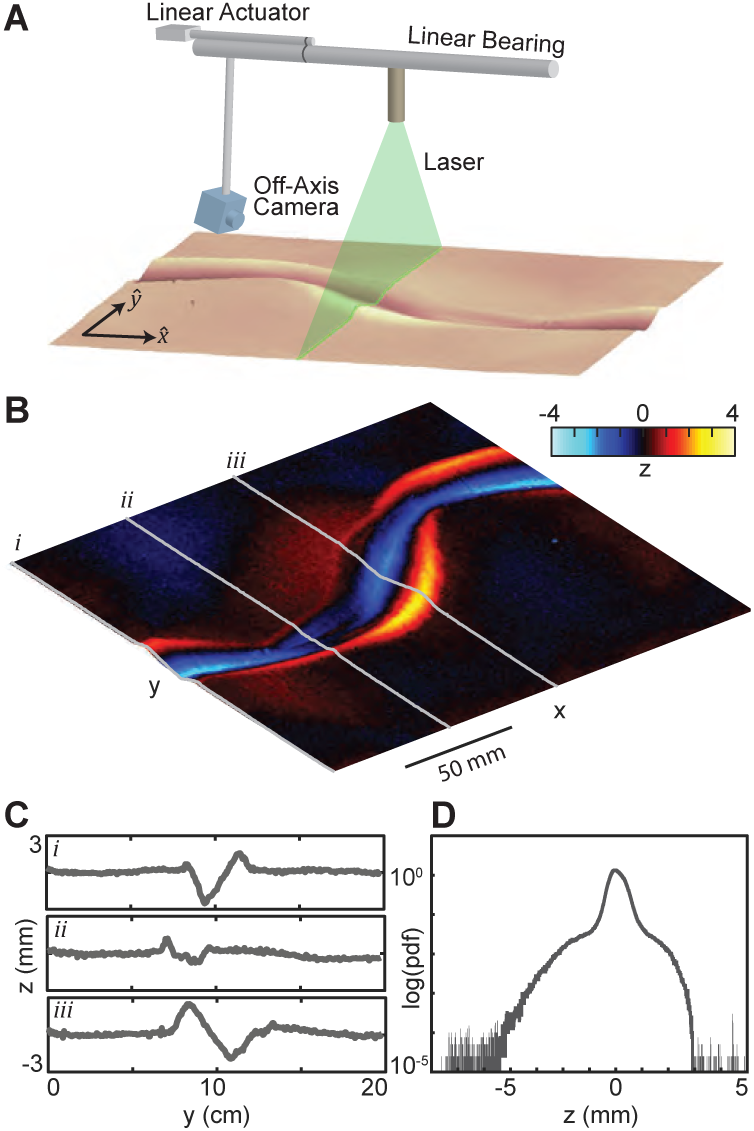
*C. occipitalis* tracks (left) Laser-line apparatus. A laser sheet spans the sand-fluidized bed. The off-axis camera detects changes in the height of the line. The linear bearing allows the laser and camera to move in x and reconstruct the snake tracks in 3D (D) Buildup of a granular pile on the front face of the intruder at y = 90o. (right) Track in the GM remaining after a trial. Warm colors indicate the piles formed rising above the free surface. The snake travelled in the direction indicated. The body intrudes beneath the free surface and yields the material, creating piles on the anterior edges of the path like those described by Mosauer and seen in the tracks observed in the field.

**FIG. S5.**
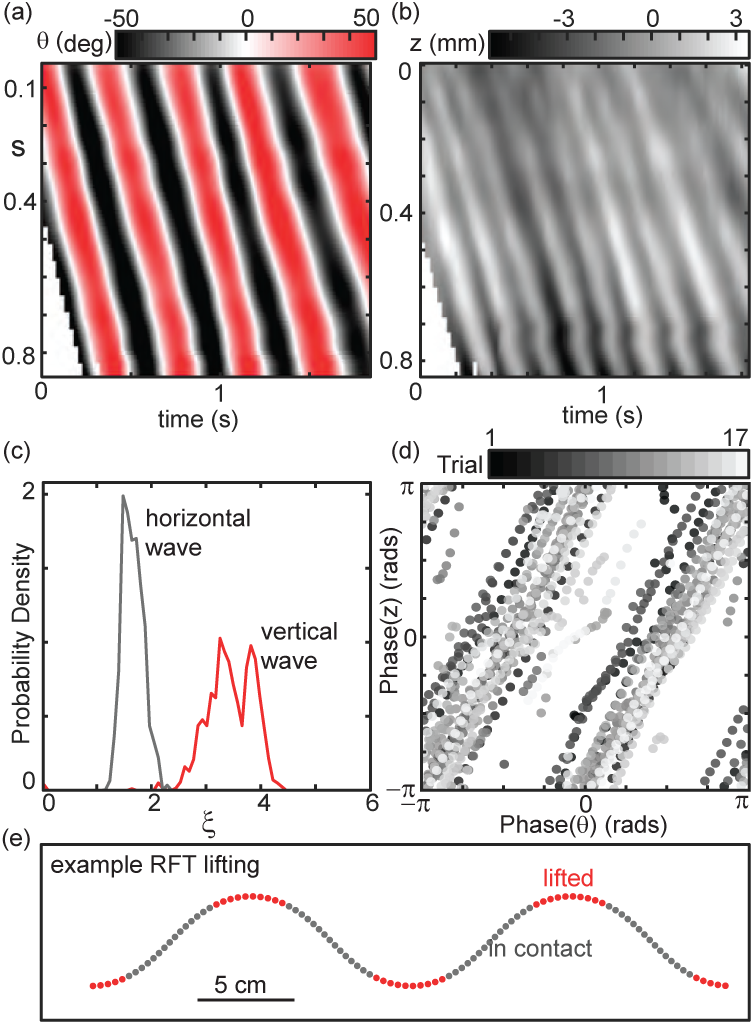
Measurement of the vertical wave of lifting. (a) Space-time plot of *θ. s* is the fractional arclength from the neck at *s* = 0 to the vent at *s* = 1. (b) Space-time plot of the vertical position of the belly of the snake measured relative to the free surface. Axes are as in (a). (c) Probability density of the wavenumber of the horizontal wave (gray) and vertical wave (red). The ratio of the median values of the two curves is 2.1, indicating that the spatial frequency of the wave of lifting is twice that of the wave in the horizontal plane. There were some outliers due to measurement error which were *ξ >* 10 that have been excluded as unphysical. (d) The phase of the wave in *z* versus the wave in *θ*. The wave in *z* is traveling at twice the frequency of that in *θ*. The result is that the peaks of the wave in *z* align with the extrema in *θ*, that is, the apexes of the horizontal wave are lifted off of the substrate. N = 3 individuals (122-11 trials, 124-3 trials, 131-3 trials) Ratio of the median values of each curve is 2.1

**FIG. S6.**
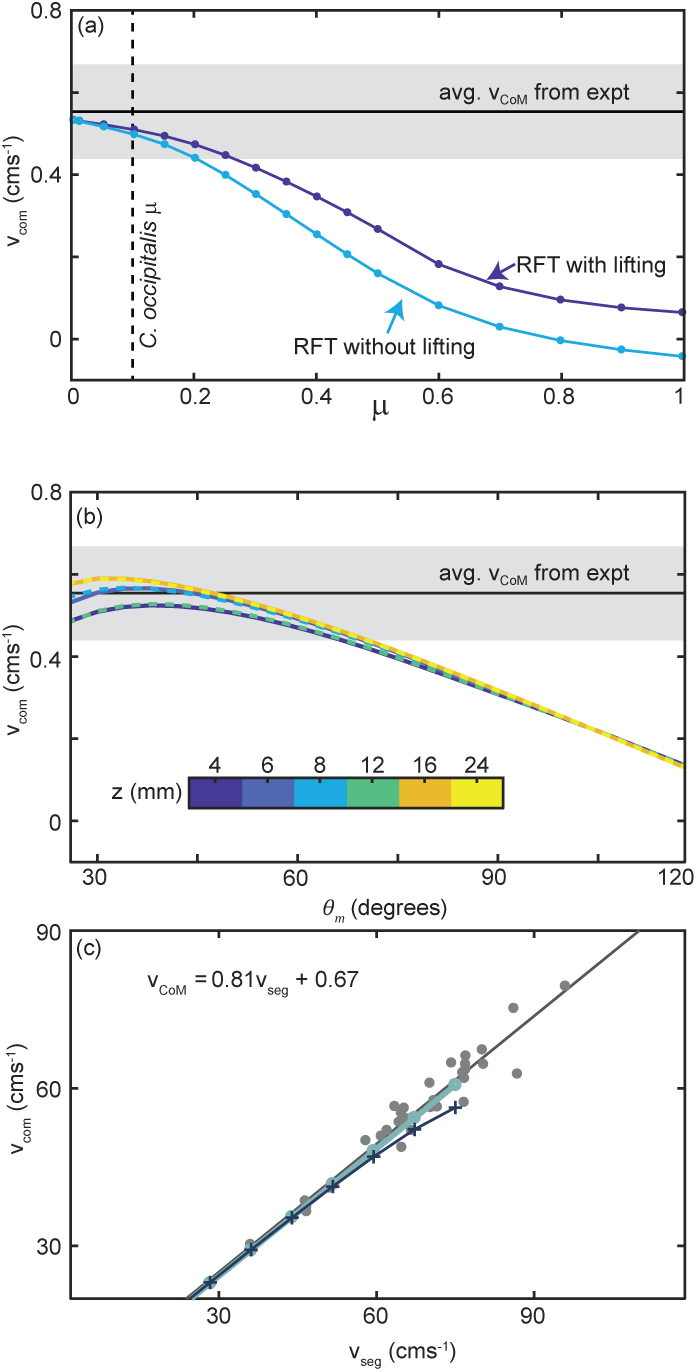
Comparison of RFT results with snake performance and impact of changing parameters. (*a*) *v*_*CoM*_ as a function of the anisotropy factor–an overall multiplier on the perpendicular forces. An anisotropy factor of two corresponds to the animal, as the low-friction scales of the snake reduces the friction by a factor of two as compared to the Aluminum plate used in the drag measurements. The blue line is RFT calculation for the snake parameters without any lifting, light blue calculated for lifting of segments which are in the top 38% of curvatures, and green dashed for in the top 73%. The black line and gray bar is average and range of *v*_*CoM*_ calculated from experiment. The RFT prediction and animal measurements come into agreement at an anisotropy factor of two, as expected. We also see that lifting does not greatly impact *v*_*CoM*_, nor does decreasing friction (thereby increasing the anisotropy factor) beyond the value achieved by the snake. (*b*) RFT calculation of *v*_*CoM*_ as a function of *θ*. The different colors indicate force relations obtained by fitting data taken at the noted depths. We find that the RFT prediction was relatively insensitive to depth which we expected given the depth independence of *K*. Further, the peak of the curves does not depend on depth. Therefore, we use in the calculations force relations measured at a depth of 8 mm.

**FIG. S7.**
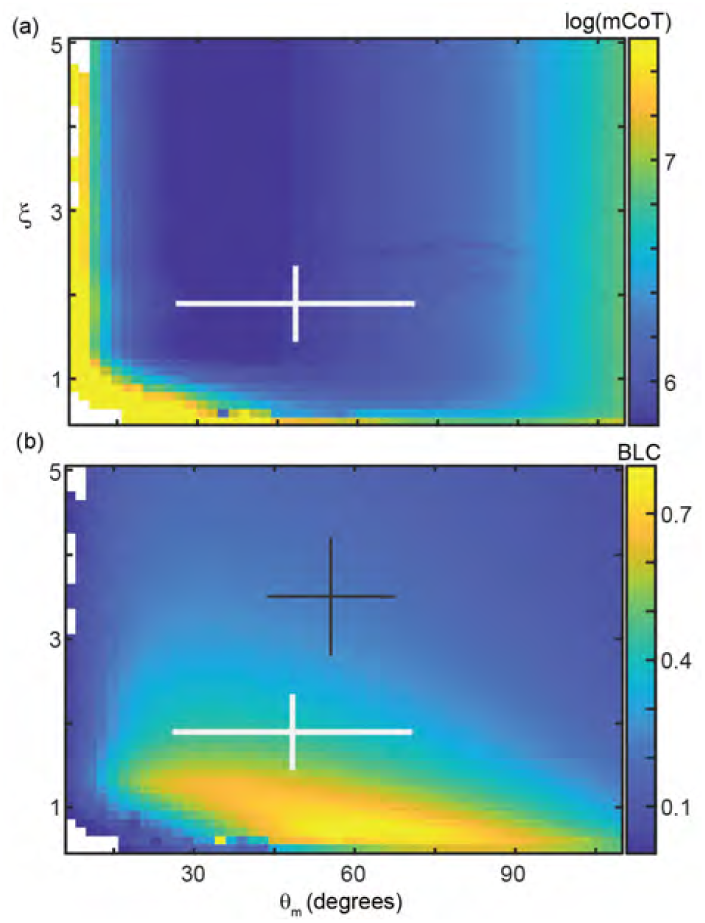
Surface RFT results for mCoT, BLC, and. *τ*_*m,RFT*_. The horizontal axis of all plots is *θ*_*m*_. Moving from left to right increases how sharply bent the body wave is. The vertical axis is *ξ*, the number of waves on the body increases from bottom to top of each plot. White crosses are average ± range of mean values measured in 30 snake trials (N=9 individuals). (a) Mechanical cost of transport, mCoT, calculated for each pair of *θ*_*m*_ and *ξ*. The color corresponds to log(mCoT). (b) Bodylengths traveled pedr cycle, BLC. Black cross is the waveform used by *C. occipitalis* when moving subsurface within the same ≈ 300 *µ*m glass particles. Values from [5].

**FIG. S8.**
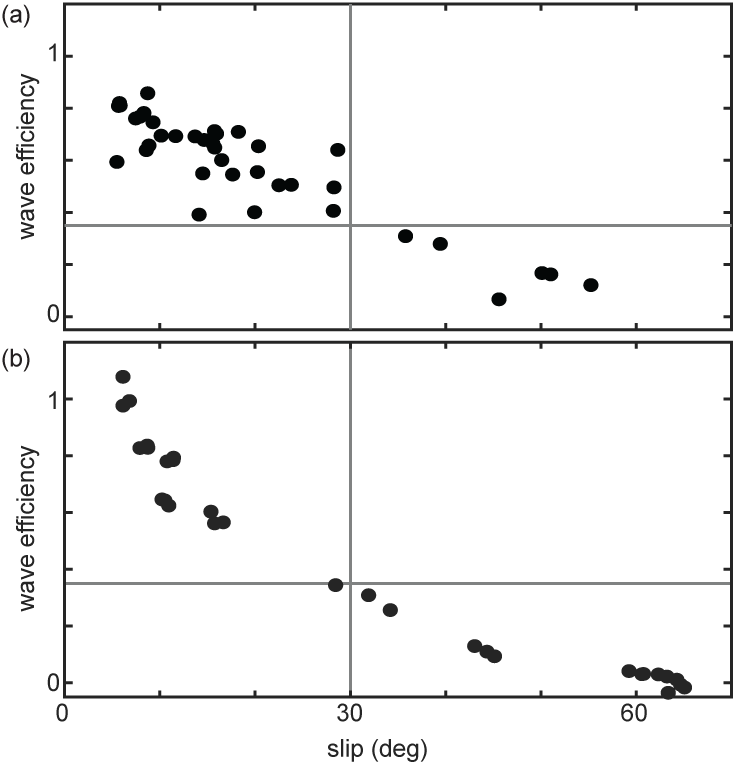
Wave efficiency versus slip. (a) Wave efficiency, average distance traveled in one undulation cycle divided by the average wavelength, as a function of slip. Black markers are from each experimental trial taken using the various snakes at the zoo on natural sand. Vertical gray line is at *β*_*s*_ = 30 deg and horizontal line is at wave efficiency of 0.35. (b) Wave efficiency, distance traveled in one undulation cycle (calculated as the total distance traveled divided by three) divided by the wavelength, as a function of slip. Black markers are from each experimental trial taken using the robot for all trials wavenumber 1, 1.2, and 1.4. Vertical gray line is at *β*_*s*_ = 30 deg and horizontal line is at wave efficiency of 0.35.

## 5. Supplementary movies

**FIG. M1.**
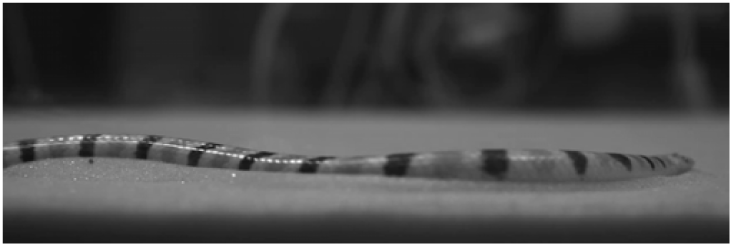
Video: Side view of *C. occipitalis* moving on GM. Video is slowed ten times.

**FIG. M2.**
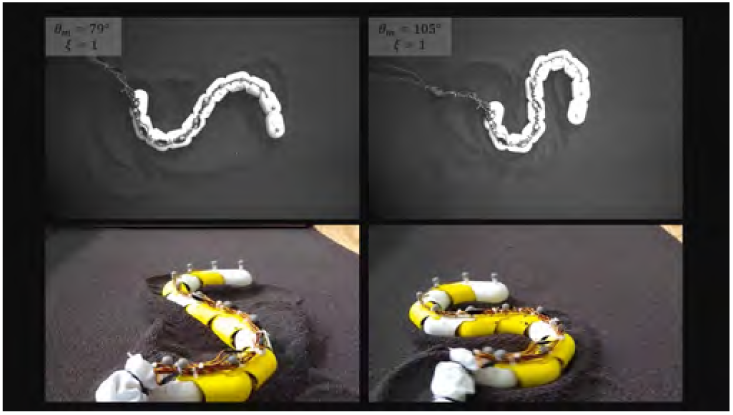
Video: Snake robot moving on poppy seeds. Three undulations Top row is overhead video, bottom row is an oblique view from behind the robot (tail is at bottom of video). Left column is an intermediate waveform (*θ*_*m*_ = 79 deg, *ξ* = 1). Right column is high attack angle (*θ*_*m*_ = 105 deg, *ξ* = 1). Waveform parameters are also displayed in the top left corner of each overhead video.

**FIG. M3.**
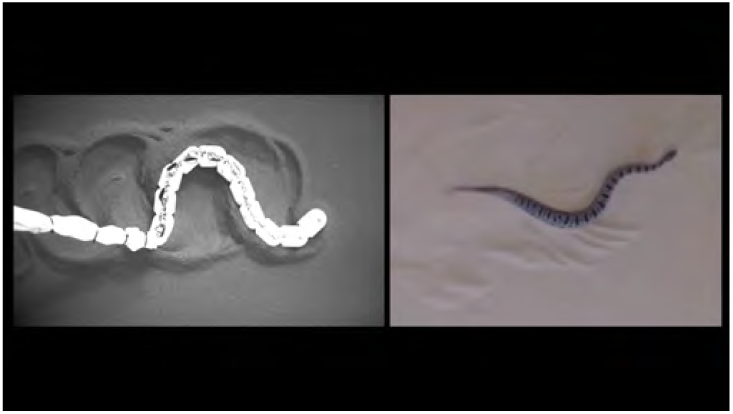
Video: Robot and snake failure comparison. To the left is the robot moving on poppy seeds using an intermediate (79 deg) attack angle with one wave on the body, to the right is the Pygmy Rattle snake, *Sistrurus miliarus* on Yuma sand. The robot video begins after the robot has completed two full undulation cycles.

**FIG. M4.**
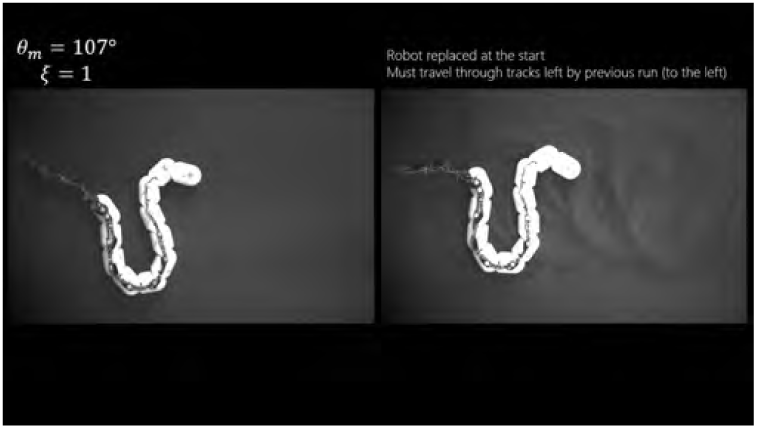
Video: Snake robot replaced in old tracks. Movie of the snake robot moving across the surface of poppy seeds using one wave on the body and a high (107 deg) attack angle. Left video shows three undulations on undisturbed poppy seeds, right is three undulations where the robot has been placed back in the track made during the trial to the left.

